# Pregnancy and Early-Life Gut Virome in the Lifelines NEXT cohort: Origin, Persistence, Influencing Factors and Health Implications

**DOI:** 10.64898/2026.03.05.709809

**Authors:** Asier Fernández-Pato, Anastasia Gulyaeva, Nataliia Kuzub, Trishla Sinha, Siobhan Brushett, Johanne E. Spreckels, Milla Brandao Gois, Archontis Goumagias, Angel Ruiz-Moreno, Antonio Pedro Camargo, Lifelines NEXT cohort study, Jingyuan Fu, Alexander Kurilshikov, Simon Roux, Sanzhima Garmaeva, Alexandra Zhernakova

**Affiliations:** Department of Genetics, University of Groningen, University Medical Center Groningen, Groningen, the Netherlands; Department of Health Sciences, University of Groningen, University Medical Center Groningen, Groningen, the Netherlands; Department of Medical Oncology, University of Groningen, University Medical Center Groningen, Groningen, the Netherlands; Department of Pediatrics, University of Groningen, University Medical Center Groningen, Groningen, the Netherlands; US Department of Energy, Joint Genome Institute, Berkeley, California, USA; Department of Biochemistry, Institute of Chemistry, University of São Paulo, São Paulo, SP, Brazil

**Keywords:** infant gut virome, pregnancy gut virome, breastmilk virome, mother-to-infant transmission, phage persistence

## Abstract

The human gut virome is a key modulator of gut microbial ecology and function, yet its role in early gut ecosystem development remains poorly understood. Here, we profiled the DNA virome from 4,523 fecal and 91 breastmilk metagenomes from 714 mother–infant pairs in the Dutch birth cohort Lifelines NEXT. This analysis generated a catalog of 31,205 unique vOTUs, with 31,019 detected in fecal and 248 in breastmilk samples, including 16,540 not previously reported in other databases. We find that the maternal virome is largely stable, in contrast to the infant virome’s rapid diversification over time. We also identify delivery and feeding modes as major drivers of infant virome developmental trajectories, with additional influences of maternal parity, infections during pregnancy, socioeconomic factors, gestational age and infant birth weight. Notably, increased viral diversity was associated with the infant developing a food allergy. Strain-level virome profiling confirmed the maternal gut as the primary source of viruses for the infant gut, with increased sharing rates in vaginally delivered infants and with breastmilk as a secondary reservoir. We demonstrate that temperate phages frequently co-transmit with their bacterial hosts and identify multiple protein families associated with anti-defense functions enriched among maternally shared viruses. Finally, we show that DNA adenine N6-methyltransferase *hin1523*, together with widely active diversity-generating retroelements, promote long-term viral persistence in the infant and maternal gut. Together, these findings establish the origin, dynamics and modulating factors of the infant gut virome, along with the genetic strategies supporting its persistence in the gut ecosystem.

## Introduction

The infant gut microbiome is established rapidly following birth and plays a crucial role in shaping host immune system maturation and metabolic function^1–4^. The gut of newborns is first colonized by pioneering microbes derived from maternal and environmental sources^5–7^. This early low diversity stage is followed by a progressive increase in microbial richness and complexity, and the community reaches a stable, adult-like configuration around 3–5 years of age^8,9^. Early gut microbiome development is influenced by maternal and birth-related factors, including mode of delivery, while later microbial maturation is shaped by infant feeding practices, medications and environmental exposures^7,10–12^.

An essential but understudied component of the human gut microbiome is the virome, comprising mostly bacteriophages, but also containing eukaryotic and archaeal viruses^13,14^. Ecological and experimental studies have demonstrated that the virome strongly shapes the composition and dynamics of gut microbial communities, highlighting the need to investigate its role in early-life gut ecosystem development^15,16^. Until recently, however, virome research had been limited by methodological and technical challenges, including difficulties in cultivating and characterizing viral isolates^17,18^. Emerging studies of the infant gut virome have now begun to reveal its potential roles in modulating the host immune system and in the development of diseases such as asthma, atopic disease and type 1 diabetes^19–21^.

Only a few longitudinal studies, with limited sample sizes, have examined the infant gut virome with paired maternal sampling^22–26^. These studies indicate that viruses can be transmitted from mothers to infants, supporting a model in which early viral colonizers are largely prophages induced from early-colonizing bacteria^27^. However, the mechanisms of transmission and the extent and functional relevance of the maternally transmitted phages remain unclear. Moreover, investigations of microbial and viral transmission have largely focused on transmission from the maternal gut to the infant gut, as technical challenges have limited metagenomic characterization of other potentially relevant microbial communities, e.g., that in breastmilk. Although still limited, existing studies indicate that breastmilk harbors diverse viral populations that can be transmitted vertically from mother to infant^25,28^. Strain-level characterization of viruses in breastmilk, alongside other environmental sources, is therefore critical to understand their contribution to infant gut colonization and early gut ecosystem development.

Beyond initial colonization, it is crucial to understand the strategies that enable viruses to persist in the gut. Several potential mechanisms of viral persistence linked to the molecular and ecological biology of viruses have been proposed, including spatial heterogeneity within the gut, enhanced adaptation to the gut environment through mucosal adherence or immune tolerance, bacterial host persistence and stop codon reassignment^29–32^. Further, previous studies have associated phage population diversity to viral persistence, suggesting that greater diversity may help in evading bacterial defense systems^33,34^. However, despite a rapid increase in the discovery of viral counter-defense mechanisms, with more than 150 new systems identified to date^35,36^, no study has evaluated their contribution to viral persistence. Likewise, a recent metagenomic analysis found that diversity-generating retroelements (DGRs) are common in human gut phages, suggesting they play a role in persistence through hypermutation of host-attachment proteins, enabling a broader host range^37^. Yet, no large-scale study has explored DGR distribution, activity and function in the early gut viral ecosystem.

Here, we aimed to provide a comprehensive view of the early gut virome assembly and maternal gut viral dynamics during pregnancy. Using 4,523 fecal metagenomes longitudinally collected from 714 mother–infant pairs in the prospective birth cohort Lifelines NEXT (LLNEXT)^38^, we generated an extensive gut viral catalog comprising over 31,000 virus operational taxonomic units (vOTUs) and investigated their associations with host health and development. We also profiled the breastmilk virome for 91 mothers to explore its contribution to infant viral colonization. By performing strain-level profiling of all vOTUs, we identified and functionally characterized potentially transmitted viruses from both maternal sources. Finally, we characterized viral anti-defense repertoires and explored DGR presence and activity to identify genetic features that support long-term viral persistence in the gut.

## Results

### LLNEXT gut viral catalog: generation and characteristics

Using fecal metagenomic samples from 714 mother–infant pairs enrolled in LLNEXT, we conducted a longitudinal analysis to characterize DNA viruses in the maternal and infant gut. Our dataset includes 1,587 maternal fecal samples collected at three pregnancy time points (12 weeks (P12), 28 weeks (P28) and birth (B)) and one postnatal time point (3 months postpartum (M3)) and 2,936 infant fecal samples collected at seven time points during the first year of life (from the second week of life (W2) up to 12 months (M12)) (**Fig. 1a**). In addition to biological samples, we analyzed 42 variables capturing clinical, perinatal, dietary, environmental and infant health and growth–related information (**Supplementary Table 1**). Comprehensive cohort characteristics are detailed in our previous study^7^.

**Fig. 1.**
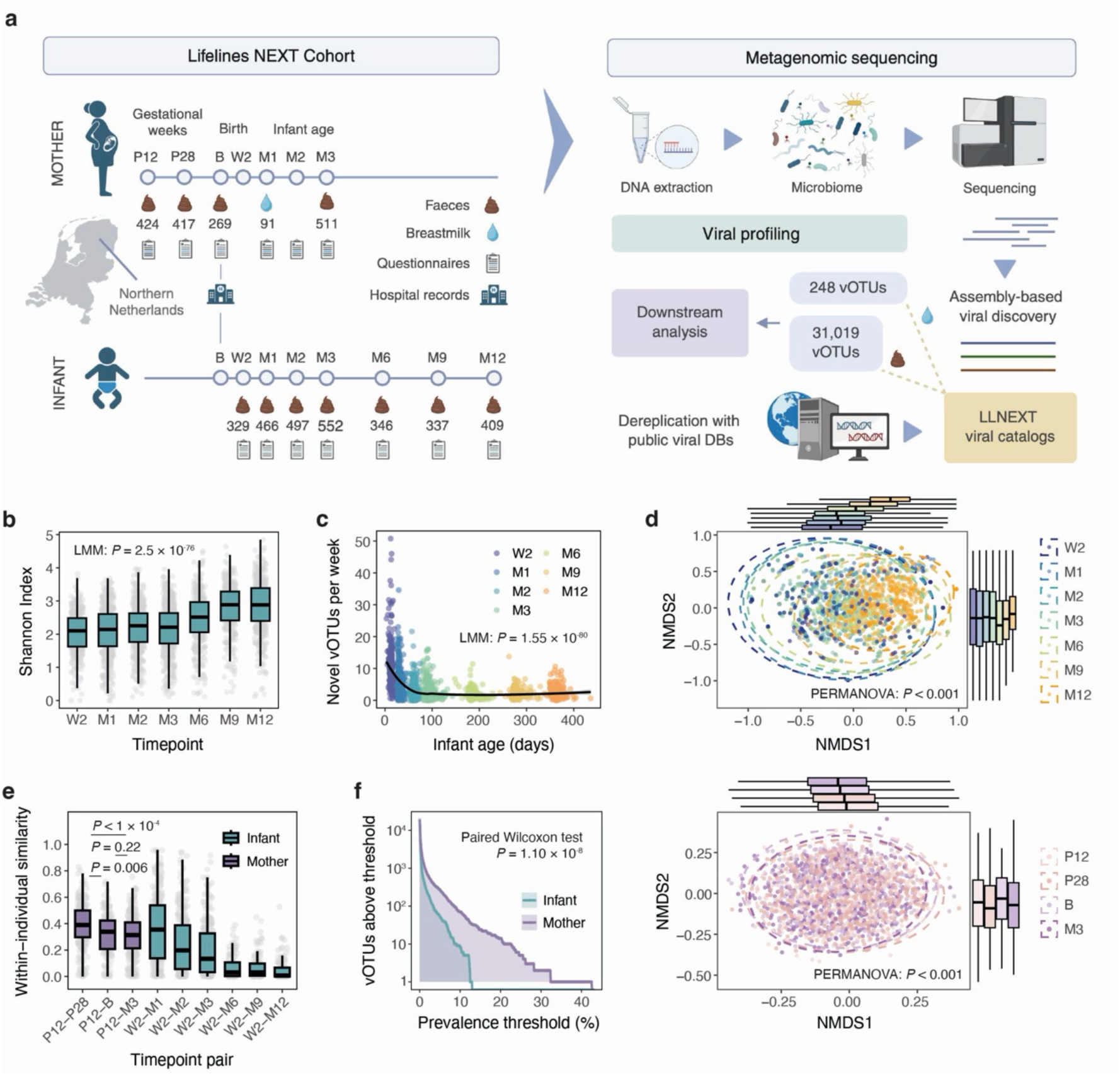
Lifelines NEXT cohort design, viral catalog generation and overview of gut virome dynamics. **(A)** Overview of the prospective mother–infant cohort Lifelines NEXT (LLNEXT) and its longitudinal sample collection. Fecal samples were collected from mothers during pregnancy (P12, P28), at birth (B) and at 3 months postpartum (M3) and from infants at week 2 (W2) and months 1, 2, 3, 6, 9 and 12 (M1‒M12). Additional breastmilk samples were collected from a subset of mothers at M1. Detailed metadata was obtained through questionnaires and hospital records. Metagenomic sequencing, followed by assembly-based viral discovery and dereplication with public viral databases, resulted in the LLNEXT gut (LLNEXT-GV) and breastmilk (LLNEXT-BMV) viral catalogs comprising 31,019 and 248 vOTUs, respectively. **(B)** Shannon diversity in infant samples over the first 12 months of life. **(C)** Number of new vOTUs detected per week of infant age. Each point represents one sample, colored by timepoint. A LOESS smoothing curve (black line) highlights the overall trend in new vOTU acquisition over time. **(D)** Non-metric multidimensional scaling (NMDS) ordination of Bray-Curtis distances estimated on vOTU abundances with a prevalence of ≥5%, illustrating viral community structure across infant (top) and maternal (bottom) samples. Each dot represents a sample. Ellipses indicate 95% confidence intervals for each timepoint. Boxplots above and to the right show the distribution of NMDS axis scores by timepoint. **(E)** Within-individual similarity of viral communities based on Bray-Curtis distances across timepoint pairs for mothers and infants, calculated by comparing each later timepoint against P12 for mothers and against W2 for infants. In all cases, boxplots show the median, IQR and 1.5 × IQR (whiskers). **(F)** Prevalence distributions of vOTUs in mothers and infants, shown as the number of vOTUs (log-scaled) above increasing prevalence thresholds. Statistical tests and P-values are reported within the corresponding panels.

Using an assembly-based viral identification framework (Methods), we identified 31,031 vOTUs across all samples. After excluding potential contaminants based on taxonomic assignment and comparison with negative controls (four vOTUs assigned to the *Sinsheimervirus phiX174* species and two prevalent infant vOTUs predicted to infect the *Phyllobacterium* genus) and RNA viruses (six vOTUs assigned to the *Riboviria* realm), we retained 31,019 vOTUs in our maternal‒infant gut DNA viral catalog (hereafter referred to as the LLNEXT-GV catalog). 16,390 of these vOTUs are unique to our dataset, showing no species-level clustering with viral genomes from public databases (**Extended Data Fig. 1a**). Quality assessment revealed that 18,172 vOTU-representative sequences (58.6%) were classified as at least high quality, with 3,637 (11.7%) vOTUs classified as complete viral genomes. The majority of vOTUs were predicted to be bacteriophages, predominantly within class *Caudoviricetes* (n = 30,340, 98.1%), with most exhibiting a temperate lifestyle (n = 19,647, 63.3%). Bacterial host prediction assigned a genus-level host to 89.1% of vOTUs, with *Bacteroides*, *Escherichia* and *Bifidobacterium* the most common predicted hosts (**Extended Data Fig. 1a**). Clustering of vOTUs into higher taxonomic ranks yielded 2,399 genus-level and 449 family-level clusters. Almost 10% of vOTUs (n = 3,090) also carried predicted viral anti-defense systems (ADS), which allow phages to overcome bacterial immunity^35,36^. The most prevalent systems were anti-restriction modification (anti-RM) (33.8%), anti-CRISPR (23.4%) and anti-Thoeris (21.5%) (**Extended Data Fig. 1b**).

Focusing on early life, we identified 16,803 vOTUs present in infant gut samples. Of these, 11,586 were not detected in maternal samples and 8,821 were absent from other human gut viral databases, representing a valuable resource for characterizing gut phage and viral communities during infancy. Notably, the ADS profiles of infant vOTUs were highly dynamic (**Extended Data Fig. 1c**), with strong enrichment of anti-CBASS (Fisher’s exact test, false discovery rate (FDR) = 5.18 × 10⁻^43^) and anti-RecBCD (Fisher’s exact test, FDR = 3.27 × 10^−55^) systems (**Supplementary Table 2**). These systems were particularly prevalent in phages targeting *Klebsiella* (FDR = 9.81 × 10^−105^, odds ratio (OR) = 76.6) and *Escherichia* (FDR = 3.29 × 10^−87^, OR = 18.1, Fisher’s exact test), respectively, reflecting how early-life bacterial abundances drive the differences observed in ADS profiles (**Extended Data Fig. 1d**).

### The gut virome is individual-specific, showing temporal stability in mothers and dynamic changes in infants

To elucidate the overall dynamics of viral communities in the infant and maternal gut, we first analyzed longitudinal changes in diversity and composition. Over the first year of life, infant gut viromes exhibited increasing Shannon diversity (β = 0.07, p = 2.49 × 10^−76^, linear mixed model (LMM)) (**Fig. 1b**) and richness (β = 1.99, p = 2.3 × 10^−120^, LMM) (**Extended Data Fig. 2a**). Notably, the rate of colonization by new vOTUs was highest at early timepoints and decreased over time (β = −0.44, p = 1.55 × 10^−80^, LMM) (**Fig. 1c**), reflecting early microbial succession followed by gradual stabilization of the infant gut virome. In contrast, maternal viromes showed stable diversity across pregnancy, with only minimal differences detected between pregnancy (P12) and postpartum (β = −0.08, p = 0.029, LMM) (**Extended Data Fig. 2b**). Compositional analysis using Bray-Curtis dissimilarity revealed significant temporal shifts in infant virome composition (PERMANOVA, R² = 0.0043, p < 0.001, 1,000 permutations) (**Figure 1D**), (**Fig. 1d**), accompanied by a significant decrease in temperate phage abundance (β = −0.97, p = 6.41 × 10^−16^, LMM) (**Extended Data Fig. 2c**). Maternal virome composition exhibited subtle but significant temporal variation, (PERMANOVA, R² = 0.0017, p < 0.001, 1,000 permutations) (**Fig. 1d**), with only minor changes in within-mother composition between early pregnancy (P12) and later timepoints (**Fig. 1e**). Overall, maternal samples showed substantially higher within-individual similarity than infants (median: 0.35, interquartile range (IQR) 0.23–0.45 vs. median: 0.098, IQR 0.02–0.29; permutation test, 10,000 permutations, p < 1 × 10^−4^) (**Fig. 1e and Extended Data Fig. 2d**). Interestingly, virome similarity also reflected biological-relatedness, with related infants (siblings and twins) showing consistently higher similarity compared to unrelated infants. We observed a similar trend for mothers, with viral profiles within and across pregnancies in the same mother (n = 18) more alike compared to those of unrelated mothers (permutation test, 10,000 permutations, *p* < 1 × 10^−4^) (**Extended Data Fig. 2e,f**). In total, 84.9% (45 of 53) of prevalent vOTUs showed significant temporal changes in infants, compared to only 14.1% (10 of 71) vOTUs in mothers (prevalence ≥ 5% in infants and ≥ 10% in mothers, log-transformed abundances) (**Supplementary Tables 3 and 4**) (**Extended Data Fig. 3a,b**). Similarly, host-level analysis showed that the abundance of phages predicted to infect 37 different bacterial genera (92.5% of prevalent phage groups by host genus) changed in infants over the first year of life, whereas only 5 phage groups (8.1%) showed significant variation in mothers (**Extended Data Fig. 3a,b**) (**Supplementary Tables 5 and 6)**. Notably, none of the host-level phage groups showed differences in abundance when analyzing only the pregnancy period (P12–B) (**Supplementary Table 7**). Both maternal and infant gut viromes were highly individual-specific, with 17,630 vOTUs (56.8%) unique to single individuals. This effect was more pronounced in the infant gut, where vOTU prevalence was consistently lower than in mothers (p = 1.10 × 10^−8^, paired Wilcoxon test) (**Fig. 1f**). Only 53 vOTUs (50 *Caudoviricetes*, 2 *Malgrandaviricetes*, 1 *Arfiviricetes*) were detected in ≥5% of infants, with no vOTUs shared across >15%, highlighting that the developing gut viral ecosystem is dynamic and individual.

### The infant gut virome is shaped by delivery mode, feeding mode and maternal factors and associated with food allergy development

In contrast to the better-characterized modulation of bacteriome assembly, the factors shaping early-life gut viral community assembly and dynamics remain poorly understood. To explore this, we assessed associations between multiple virome features (richness, alpha diversity, viral abundances and the relative abundance of temperate phages) and a comprehensive set of 42 phenotypes (maternal health and dietary patterns, birth characteristics, perinatal exposures and early-life clinical outcomes) (**Supplementary Table 1**). Given the known strong influence of delivery and feeding modes on gut microbial colonization^7,10,11,39^, we performed all analyses both unadjusted and adjusted for these covariates. Indeed, viral diversity, but not viral richness, was positively associated with formula feeding (p = 5.42 × 10^−3^ and p = 0.086, respectively; LMM), while both richness and diversity were positively associated with vaginal delivery (p = 9.52 × 10^−3^ and p = 7.84 × 10^−3^, respectively; LMM) at nominal significance (**Fig. 2a and Extended Data Fig. 4a**) (**Supplementary Table 8**). Importantly, pre-labor Cesarean section (CS) was associated with greater viral diversity than CS performed after onset of labor (p = 6.81 × 10^−3^, LMM). Several maternal factors were also nominally associated with infant viral richness and diversity (p < 0.05, LMM adjusted for feeding and delivery mode), including pregnancy infections (vaginal fungal infection, flu/respiratory tract infection, gastroenteritis, urinary tract infection), pre-pregnancy smoking history, maternal parity (number of previous births) and income, specific dietary patterns and delivery assisted with a vacuum pump, suggesting that the maternal and early environment contribute substantially to virome assembly (**Supplementary Table 9**). Importantly, infant food allergy in the first year of life, defined as any parent-reported food allergy or adverse food reaction, was associated with increased viral diversity (p = 5.32 × 10^−3^), and this remained nominally significant even after adjusting for bacterial alpha diversity (**Fig. 2b**) (**Supplementary Table 10**). In addition, maternal income, which likely reflects socioeconomic and lifestyle differences, and vaginal fungal infections were significantly associated with higher infant viral diversity after multiple testing correction and adjustment for bacterial diversity (FDR < 0.05, LMM) (**Extended Data Fig. 4b**) (**Supplementary Table 10**).

**Fig. 2.**
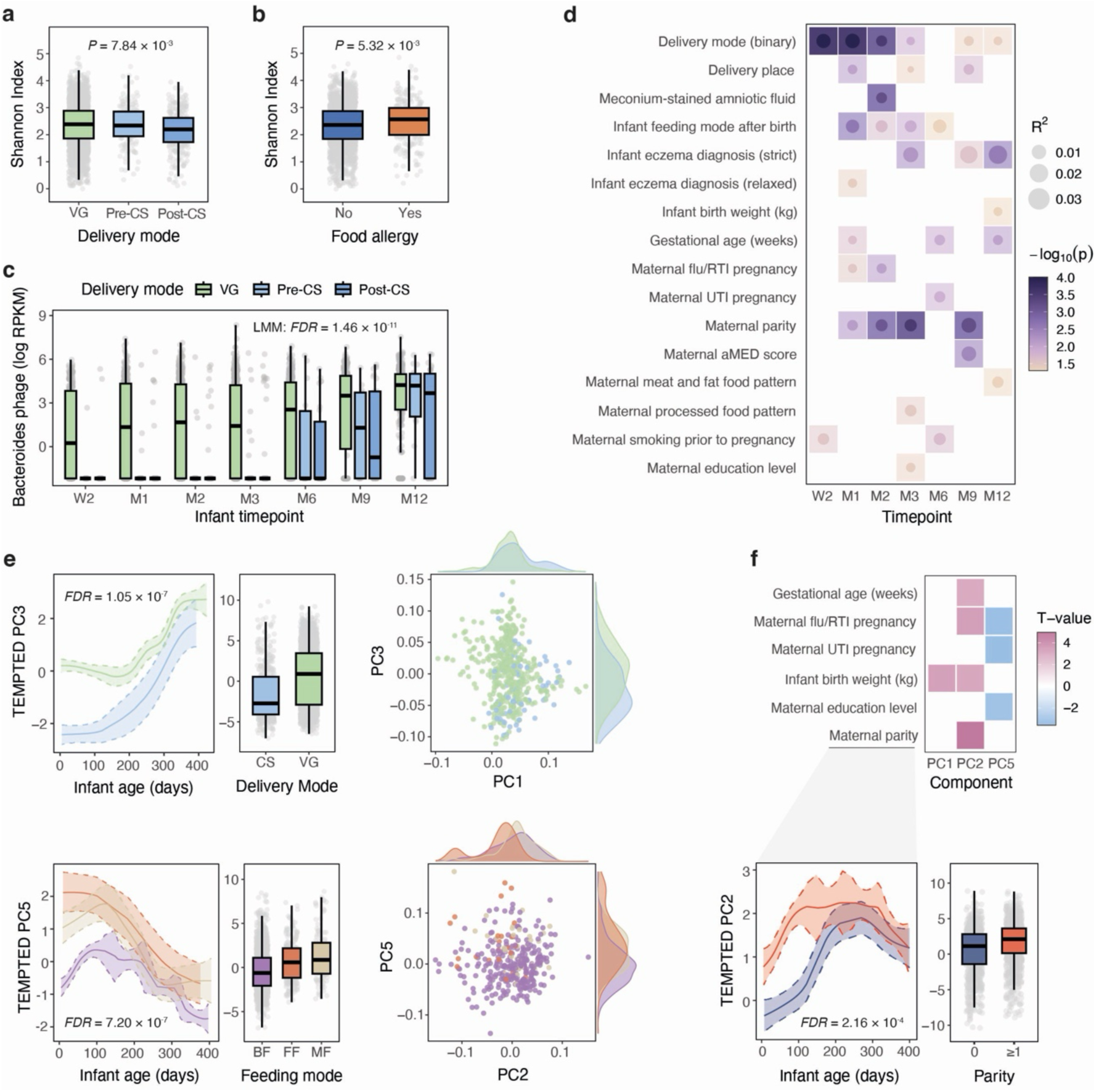
Factors shaping the infant gut virome and its association with infant health outcomes. **(A–B)** Associations between Shannon diversity and **(A)** delivery mode (VG: vaginal delivery, Pre-CS: Cesarean section before labor, Post-CS: Cesarean section after labor) and **(B)** infant food allergy status. **(C)** Abundance of Bacteroides phages (log RPKM) across infant timepoints, stratified by delivery mode. **(D)** Associations between infant virome composition (Bray-Curtis dissimilarities) and maternal, birth and early-life phenotypes per timepoint (PERMANOVA, 10,000 permutations). Dot size indicates variance explained (R²). Color indicates −log10(p-value). Only phenotypes with significant associations are shown (p < 0.05). **(E)** Longitudinal virome trajectories derived using TEMPTED for delivery mode (top) and feeding mode (bottom). Line plots (left) display TEMPTED component values across infant age (days of life), with shaded areas representing standard error. The most significant components associated with delivery mode (PC3) and feeding mode (PC5) are shown. Boxplots (middle) show the overall distributions of these components. Scatterplots (right) illustrate subject-level loadings for the two components most associated with each phenotype. **(F)** Associations between maternal, birth and early-life factors and TEMPTED components, adjusted for feeding and delivery mode. Only phenotypes with significant associations are shown (FDR < 0.05). Color indicates the direction and magnitude of the association (t-value). Below, a line plot and boxplots illustrate longitudinal virome trajectories based on TEMPTED component 2, stratified by maternal parity (no previous births vs. at least one previous birth). In all cases, boxplots show median, IQR and 1.5 × IQR whiskers. P-values and statistical tests are shown accordingly in the figure panels.

We also explored associations with the abundances of vOTUs grouped by their predicted bacterial hosts and human phenotypes (**Extended Data Fig. 4c**). CS delivery was strongly associated with reduced abundances of phages predicted to infect *Bacteroides*, consistent with known depletion of this genus in CS-born infants^40^ (FDR = 1.46 × 10^−11^, LMM) (**Fig. 2c**) (**Supplementary Table 11**). *Parabacteroides* phages were also decreased in the CS-born group, whereas phages predicted to infect *Enterococcus*, *Enterobacter*, *Klebsiella* and *Flavonifractor* were significantly less abundant in vaginally born infants. Notably, the association of *Enterobacter* phages with delivery mode remained significant even after adjusting for bacterial host genus abundance (β = 0.47, FDR = 5.11 × 10^−3^, LMM) (**Supplementary Table 12**), indicating both host-dependent and host-independent mechanisms of virome modulation. Infant feeding mode was also associated with phage abundances, with exclusively formula-fed infants showing higher abundances of *Akkermansia* phages (β = 0.56, FDR = 8.64 × 10^−3^) (**Supplementary Table 11**). After accounting for delivery and feeding mode, the only remaining significant associations were lower *Clostridium* phage abundance (β = −0.50, FDR = 1.09 × 10^−5^) and higher *Bifidobacterium* phage abundance (β = 0.83, FDR = 7.4 × 10^−3^) in infants of mothers with previous births, potentially reflecting the effect of siblings on the infant virome (**Extended Data Fig. 4c**) (**Supplementary Table 13**).

Analysis of factors driving viral composition per infant time point revealed that multiple phenotypes contribute to the interindividual variation, including maternal socioeconomic, dietary and lifestyle factors, and infections during pregnancy; delivery-related characteristics; and infant feeding, growth, and clinical outcomes such as eczema (PERMANOVA, 10,000 iterations, p < 0.05) (**Fig. 2d**) (**Supplementary Tables 14 and 15**). To assess phenotypes associated with longitudinal variation in the infant virome, we applied TEMPTED^41^, a dimensionality reduction method for multivariate longitudinal data, to a subset of infants with samples taken at three or more time points (n = 473). The TEMPTED analyses were performed using the first five principal components (PCs), which together explained 20.2% of the total variation. Association with infant and maternal phenotypes revealed delivery mode and feeding mode as the most significant factors shaping infant virome trajectory (FDR < 0.05, LMM) (**Fig. 2e**) (**Supplementary Table 16**). After accounting for these effects, several additional maternal and delivery-related factors also contributed to interindividual variation in virome dynamics. Among these factors, maternal parity showed the strongest association and was positively associated with PC2, highlighting sibling exposure as an important determinant of longitudinal infant virome dynamics independent of delivery and feeding mode (FDR = 2.16 × 10^−4^, LMM). Additional contributors included infant birth weight and gestational age, as well as maternal education level and infections during pregnancy, each of which was significantly associated with virome trajectories over time (FDR < 0.05, LMM) (**Fig. 2f**) (**Supplementary Table 17**).

### Maternal gut and breastmilk viromes contribute to early colonization of the infant gut

Recent studies have indicated that the maternal gut microbiome plays a key role in seeding the early microbial communities of the infant gut^7,42,43^. While a few studies have observed maternal‒infant sharing of gut viruses, comprehensive characterization of the maternal sources, transmission mechanisms and the functional roles of these potentially transmitted viruses in the developing infant gut remains limited. To address this gap, we first investigated the extent to which maternal gut viral strains were established in the infant gut. For this, we compared the strain profiles of all viruses detected at vOTU level in at least one maternal and one infant fecal metagenome. Strain-level profiling yielded 193,682 pairwise comparisons of 1,997 vOTUs, including 4,590 comparisons across 432 mother–infant pairs (**Supplementary Table 18**). Compared to unrelated pairs, related pairs harbored viral strains with higher similarity (one-sided permutation test, 10,000 permutations, p < 1 × 10^−4^) (**Fig. 3a**) and exhibited a higher proportion of strain-sharing events (p < 2.2 × 10^−16^, chi-squared test) (**Fig. 3b**). To evaluate whether viral strain profiles could be used to identify mother–infant pairs, we trained logistic regression models based on strain-similarity values (popANI) of maternal samples (any timepoint) versus each infant timepoint. These models accurately predicted related pairs, with predictions based on all infant timepoints yielding an AUC > 0.85 (**Fig. 3c**). Over the first year of life, the proportion of families in which any strain-sharing was detected increased with time (W2 vs M12: β = 1.84, p = 3.92 × 10^−13^, generalized linear mixed models (GLMM)) (**Extended Data Fig. 5a**), indicative of ongoing horizontal viral transmission between mothers and infants. However, the fraction of infant gut vOTUs shared with mothers (sharing rate) decreased over time (β = −0.026 per month, p = 6.99 × 10^−6^, GLMM) (**Fig. 3d**), suggesting that maternal viruses continue to contribute but their influence is gradually diluted by other viral sources. Importantly, delivery mode strongly influenced viral-sharing. Families with vaginal deliveries showed a higher proportion of strain-sharing events than those with infants born by CS (*p* = 1.72 × 10^−4^, chi-squared test) (**Extended Data Fig. 5b**), with strain-sharing occurring across more families (β = 1.91, p = 1.65 × 10^−7^, GLMM) (**Fig. 3e**) and at higher rate (β = 0.36, p = 0.03, GLMM) (**Extended Data Fig. 5c**) among vaginally delivered pairs. In contrast, whether infants were born at home or in the hospital did not influence viral-sharing after accounting for the mode of delivery.

**Fig. 3.**
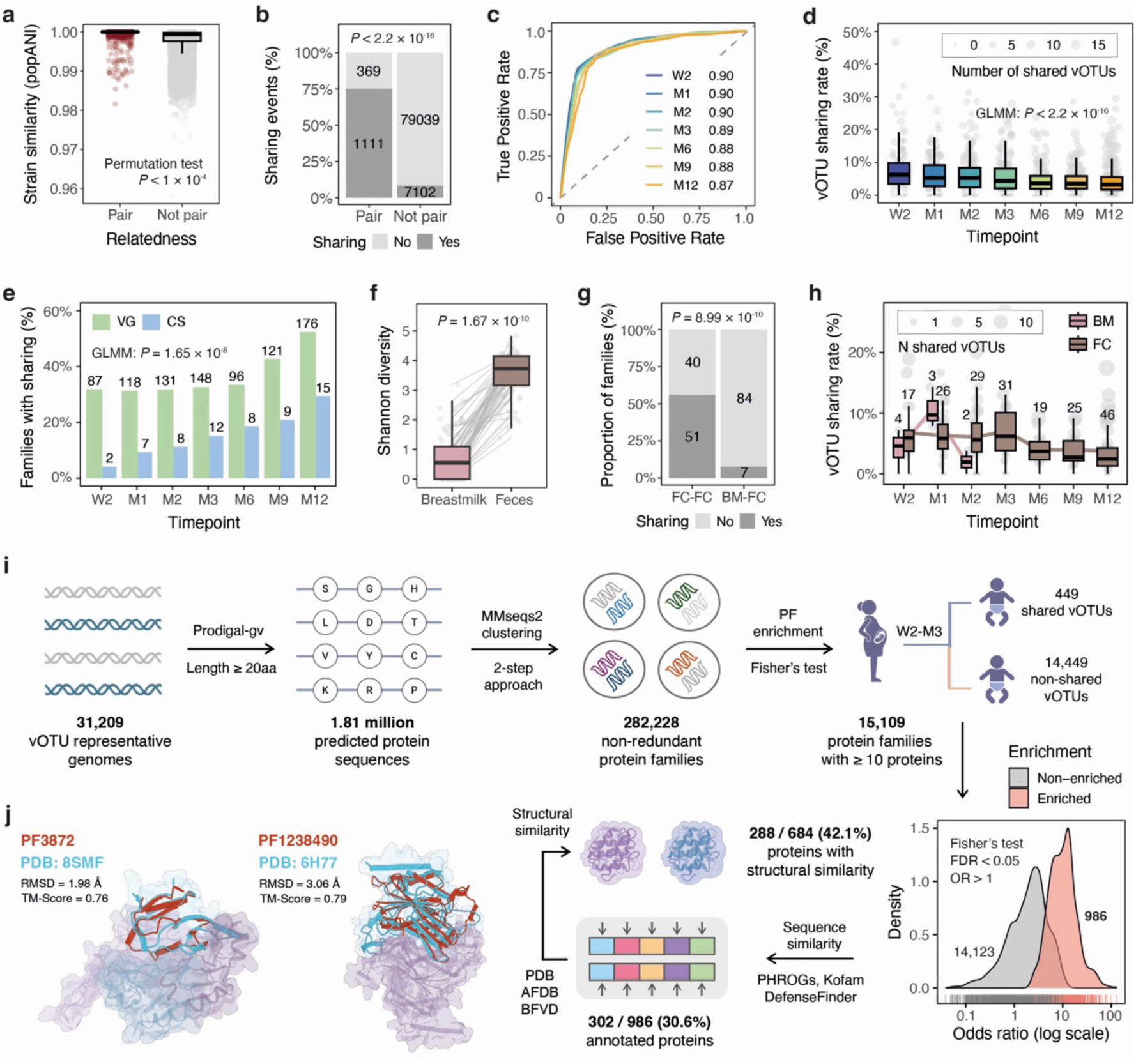
Maternal gut–infant and breastmilk–infant viral strain sharing and protein families enriched among maternally shared viruses. **A)** Viral strain-similarity (popANI) comparison between related (dark red) and unrelated (gray) mother–infant pairs. **(B)** Proportion of viral strain-sharing events in related (left) and unrelated (right) pairs. Strain-sharing events are defined as strain-level comparisons with popANI >99.999%. **(C)** ROC curves assessing the ability of viral strain profiles at each infant timepoint to distinguish mother–infant pairs, based on logistic regression models. **(D)** vOTU sharing rate (number of shared vOTUs/total number of infant vOTUs) across infant timepoints. Boxplots show the median, IQR and 1.5 × IQR whiskers. Individual gray points represent samples. Point size indicates the number of shared vOTUs. **(E)** Proportion of families with viral strain-sharing across infant timepoints, stratified by delivery mode (VG: vaginal, CS: C-section). The total number of families with strain-sharing is indicated above each bar. **(F)** Comparison of viral diversity (Shannon index) between breastmilk samples collected at M1 and fecal samples collected at B from the same mothers. **(G)** Proportion of mother–infant pairs (n = 91) in which viral strain-sharing events were detected via maternal gut–to–infant gut sharing (FC-FC, n = 51/91) or via the breastmilk–infant gut route (BM-FC, n = 7/91). **(H)** Comparison of vOTU-sharing rates across infant timepoints for maternal gut–to–infant gut versus breastmilk-mediated transmission routes. The number of mother–infant pairs with detected strain-sharing events is shown above each boxplot. **(I)** Schematic overview of the pipeline used for viral protein clustering, identification of protein families (PFs) enriched among maternally shared vOTUs and sequence- and structural-similarity–based annotation of the enriched PFs. **(J)** Comparison of the predicted structures of PF3872 and PF1238490 with PDB entries 8SMF and 6H77, respectively, using Foldseek. P-values and statistical tests are shown accordingly in the figure panels.

To investigate potential contributions of viral strains from other maternal sources, we deeply sequenced breastmilk metagenomes (40 GB per sample) from 91 mothers who breastfed their infants, with samples collected 1 month postpartum and processed using the same pipeline described previously (Methods). In total, we identified 250 vOTUs in milk samples, including 152 vOTUs with no other genomes in public databases or the LLNEXT-GV catalog (**Extended Data Fig. 5d**). Similar to the gut samples, the majority of vOTUs belonged to class *Caudoviricetes*. Among these, we detected two vOTUs predicted to infect *Microbacterium* in >30% of breastmilk samples, likely reflecting the presence of *Microbacterium* as a potential contaminant, as noted in prior microbiome studies^44^. After excluding these two vOTUs (LLNEXT-BMV catalog, n = 248), a predicted *Staphylococcus* temperate phage detected in 10 metagenomes was the most prevalent vOTU. Interestingly, one vOTU identified as human cytomegalovirus was detected in the breastmilk of three mothers. Overall, the viral diversity in breastmilk samples was significantly lower than that observed in fecal samples from the same mothers at birth (p = 1.67 × 10^−10^, Wilcoxon paired test) (**Fig. 3f**), consistent with the lower bacterial diversity reported in milk compared to gut^45^. Strain-level analysis of the 26 *Caudoviricetes* vOTUs for which strain profiles could be compared in at least one mother‒infant pair (Methods, **Supplementary Table 19**) revealed substantially lower viral strain-sharing via breastmilk than via the maternal gut–to–infant gut route (McNemar’s test, p = 5.42 × 10^−10^). Across all pairs, only 7 out of 91 showed any sharing between breastmilk and the infant gut, in contrast to 51 pairs showing gut-to-gut sharing (**Fig. 3g**). Although the limited number of breastmilk‒to–infant gut sharing events (n = 12) prevented statistical analysis of temporal trends, all the sharing events we observed occurred in infant gut samples collected at W2–M2, coinciding with active breastfeeding. For these pairs, strain-sharing rates at early timepoints were similar to those observed in gut‒gut comparisons (**Fig. 3h**). Notably, none of the vOTUs shared via the maternal gut were detected as shared with breastmilk, suggesting minimal overlap in their strain-level contribution to the infant gut. In fact, most of the vOTUs found in breastmilk were predicted to infect *Acinetobacter* (n = 41), *Streptococcus* (n = 40) and *Staphylococcus* (n = 28) (**Extended Data Fig. 5d**), all common members of the human milk microbiota that are only rarely detected in the human gut^45,46^.

### Maternal‒infant sharing of temperate viruses via co-transmission with bacterial hosts

Next, we investigated the mechanisms underlying maternal‒infant gut viral strain-sharing. First, we observed a positive association between temperate lifestyle and viral strain-sharing (β = 0.43, p = 6.35 × 10^−5^, generalized linear model (GLM)) after adjusting for the maternal mean abundance of each vOTU, suggesting that lysogeny may play a role in mother-to-infant transmission (**Extended Data Fig. 6a**). To test whether maternal-to-infant phage transfer primarily occurs via prophages integrated into transmitted bacterial strains, we mapped potentially transmitted temperate phage genomes (n = 435) to bacterial metagenome-assembled genomes (MAGs) reconstructed from the maternal and infant samples with detected strain-sharing (26,106 MAGs with at least medium-quality generated from 1,265 samples) (**Extended Data Fig. 6b**). 278 vOTUs (63.9%) mapped to at least one MAG (≥ 75% genome coverage, ≥ 95 % sequence identity), and 207 (76.4% of the 271 vOTUs with genus-level host predictions) showed concordance with the host prediction at the genus level. Notably, 38 of 64 discordant host assignments (59.4%) involved vOTUs predicted to infect *Bacteroides* that actually mapped to *Phocaeicola* MAGs, consistent with the recent *Bacteroides*–*Phocaeicola* taxonomic reclassification^47^. A subset of vOTUs aligned to multiple MAGs spanning different species (n = 44) or genera (n = 33), indicating that some phages may have broad host ranges (**Supplementary Table 20**), as also recently reported^48,49^. Among all potential sharing events (n = 670), we identified 136 cases (20.3%) where the same viral genome matched MAGs in both members of a mother–infant pair, distributed across all infant timepoints (**Extended Data Fig. 6c**). Detection of these vOTUs in pairs was also significantly enriched compared to randomly paired mother–infant samples (permutation test, p < 1 × 10^−4^) (**Extended Data Fig. 6d**). To assess whether bacteria act as mediators of phage transmission, we evaluated the whole-genome similarity of maternal–infant MAG pairs and found that 111 potential sharing events (81.6%) involved MAGs classified as the same strain (average nucleotide identity (ANI) > 99.9%), predominantly belonging to *Bacteroides uniformis*, *Phocaeicola dorei* and *Akkermansia muciniphila* (**Extended Data Fig. 6e**). Together, these findings provide strong evidence that temperate phages are often transmitted from mothers to infants via their bacterial hosts (**Supplementary Table 21**).

### Protein family‒based analysis suggests that maternally shared viruses have novel functional capacities in the infant gut

We next aimed to understand the functional relevance of viruses of maternal origin (gut and breastmilk) to the developing infant gut ecosystem in the first three months of life (W2‒M3). For this, we clustered predicted viral proteins from all viral genomes (n = 1.81 million proteins) into 282,228 non-redundant protein families (PFs) using a two-step clustering approach (Methods) (**Fig. 3i**). We then assessed PF enrichment by comparing their presence among maternally shared vOTUs (n = 449) to those found in mothers but not shared with their infants (n = 14,449). Among PFs represented by ≥ 10 proteins (n = 15,109), we identified 986 PFs that were significantly over-represented in shared vOTUs (Fisher’s exact test, OR > 1, FDR < 0.05) (**Fig. 3i**) (**Supplementary Table 22**). HMM-based annotation of representative proteins from enriched families against the PHROGs^50^, KOfam^51^ and AntiDefenseFinder^52^ databases showed that, while 625 PFs (63.4%) had a significant match to a protein profile, only 302 PFs (30.6%) matched a profile with a previously characterized function. The majority of the functionally annotated PFs were predicted to be involved in DNA, RNA and nucleotide metabolism or to have structural roles, including tail and head proteins (**Supplementary Table 22**). Notably, four enriched PFs were annotated as viral ADS, including two anti-RM PFs (hin1523 and ardc), one anti-CRISPR PF (acriia9) and one anti-Thoeris PF (acriia7).

To investigate the functional potential of uncharacterized PFs, we used multiple sequence alignments (MSAs) of unannotated clusters (n = 684) to predict a 3D structure using ColabFold^53^. This yielded 682 predicted 3D models, including high-confidence predictions (average pLDDT ≥ 70) for 635 PFs (93.1%). To infer potential functions, the predicted protein structures were searched against large-scale structural databases, including the Protein Data Bank^54^ (PDB), the AlphaFold Database^55^ (AFDB) and the Big Fantastic Virus Database^56^ (BFVD), using Foldseek^57^. The predicted structures of 288 PFs (42.1% of unannotated PFs) showed significant similarity (qtm-score > 0.7, qcov ≥ 0.7, prob ≥ 0.9) to at least one known protein structure, including 149 PFs for which no sequence-based matches were detected.

Functional inference based on structural similarity revealed several potential viral defense functions among the previously uncharacterized proteins. For instance, PF3872 showed a high-confidence structural match to the Thoeris anti-defense protein Tad2 (PDB ID = 8SMF, prob = 0.949, qTM-score = 0.76) and PF1238490 matched a Ubiquitin-like modifier-activating enzyme 5 (PDB ID 6H77, prob = 0.975, qTM-score = 0.79), potentially representing a phage homolog of bacterial ubiquitination-like proteins involved in antiphage defense^58,59^ (**Fig. 3j**). Additionally, two protein families, PF44410 (PDB ID: 4D3H, prob = 0.956, qTM-score = 0.79) and PF1562182 (PDB ID: 4D3H, prob = 0.912, qTM-score = 0.74), matched the bacterial PII-like protein PstA, consistent with recently identified phage-encoded PII-like proteins (SAM-AMP lyases) that can function in anti-CRISPR or anti-RM defense^60,61^. The full list of structural matches across all databases is provided in **Supplementary Tables 23-25**. Overall, our analysis identified multiple PFs enriched among maternally shared viruses, including metabolic, structural and other accessory functions, such as viral anti-defense roles, suggesting a maternal contribution of distinct viral genomic features to the infant gut ecosystem.

### The maternal gut shows high levels of viral persistence, with anti-RM methyltransferase *hin1523* strongly associated with persistence in mothers and infants

While maternal transmission shapes the initial seeding of the infant virome, little is known about the mechanisms that determine the persistence of gut viruses post-colonization. How these persistence patterns vary across individuals and life stages also remains unclear. To explore this, we first defined persistent vOTUs as those detected in at least three timepoints of the same infant, including at least one early (W2, M1, M2, M3) and one late (M6, M9, M12) timepoint. In mothers, persistent vOTUs were defined as those detected in at least one pregnancy (P12, P28, B) and a post-pregnancy (M3) timepoint of the same mother (Methods) (**Fig. 4a**). We identified 2,238 persistent vOTUs in infants, representing 13.32% of infant vOTUs (**Extended Data Fig. 7a**). Notably, these persistent vOTUs were typically individual-specific, i.e., persistent in only a single infant (median = 1 infant, range = 1‒40). In mothers, we found 7,395 persistent vOTUs (38.05%) (**Extended Data Fig. 7a**), again showing a strong individual-specific pattern (median = 1, range = 1–114). However, 65 vOTUs in infants and 307 vOTUs in mothers persisted across ≥ 10 individuals.

**Fig. 4.**
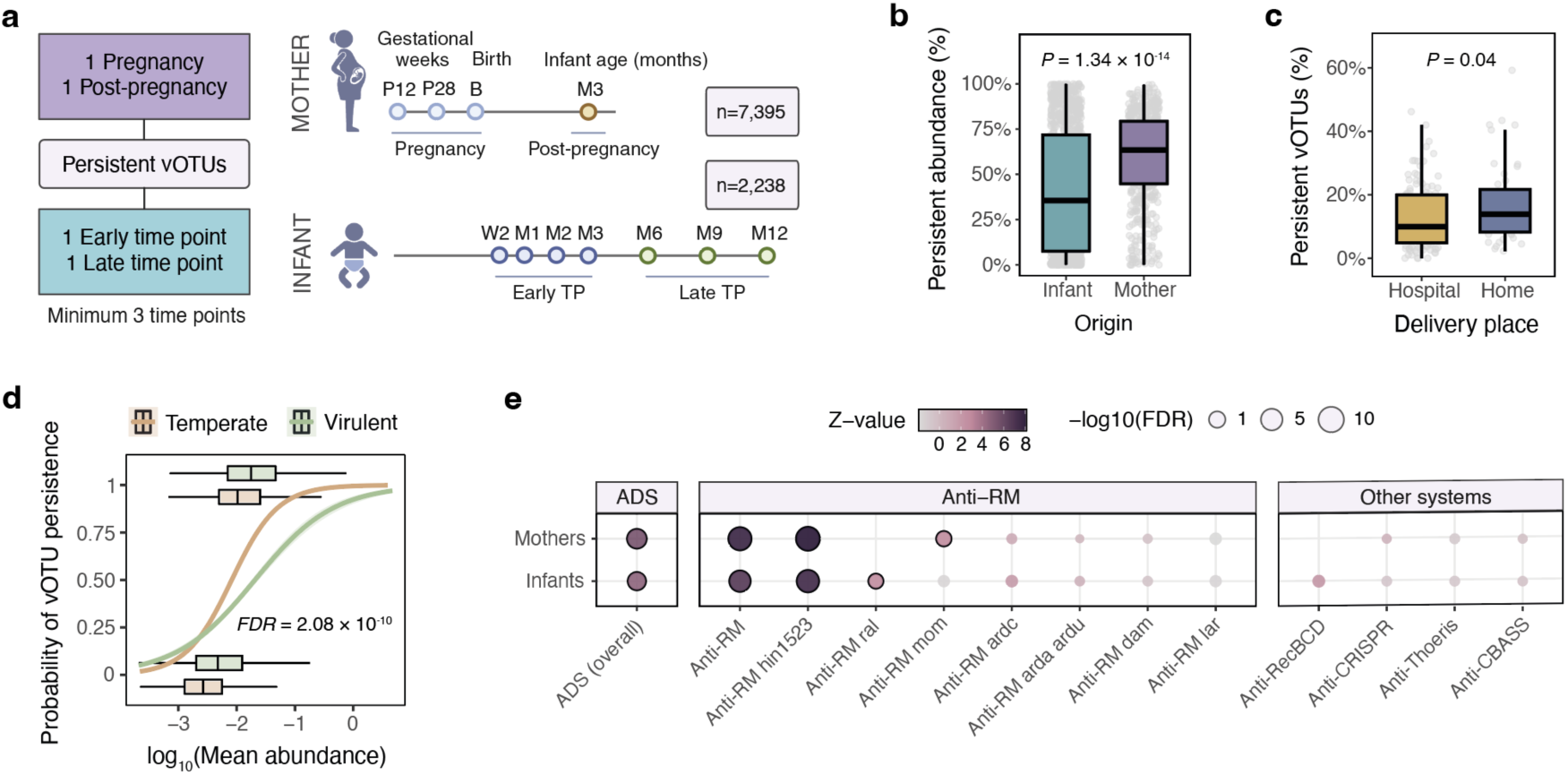
Viral persistence and associated genetic features. **(A)** Overview of the criteria used to identify persistent vOTUs. In mothers, a vOTU was defined as persistent if present in at least one pregnancy timepoint (P12, P28 or B) and one post-pregnancy timepoint (M3) in the same individual (n = 7,395). In infants, a vOTU was considered persistent if detected in at least three timepoints from the same infant, including at least one early (W2, M1, M2 or M3) and one late (M6, M9 or M12) sampling point (n = 2,238). **(B)** Comparison of the relative abundance of persistent vOTUs between mothers and infants. **(C)** Proportion of persistent vOTUs according to delivery place (hospital vs home). Boxplots show median, IQR and 1.5 × IQR whiskers. **(D)** Relationship between vOTU mean abundance in mothers and the probability of persistence, shown separately for temperate and virulent lifestyles. Logistic regression curves show the modeled probability of persistence across the range of log10-transformed mean abundances, with shaded areas representing 95% confidence intervals. Boxplots illustrate the distribution of mean abundances for persistent (top) and non-persistent (bottom) vOTUs within each lifestyle category. **(E)** Associations between viral anti-defense systems (ADS) and persistence. Each point represents the effect of the presence of a given ADS on the number of individuals in which a vOTU is persistent, estimated using a zero-inflated negative binomial GLMM adjusted for mean abundance and including bacterial host genus as a random effect. Color reflects the z-value of the system coefficient, indicating the direction and strength of the association, while point size represents statistical significance (–log10 FDR). Points with FDR < 0.05 are highlighted with a black border. Results are shown separately for mothers and infants, with anti-restriction modification (anti-RM) systems further divided into specific system subtypes. Anti-RM ral (n = 12) and anti-RecBDC (n = 27) were excluded from the association analysis in mothers due to their detection in low numbers.

To ensure an accurate assessment of persistence, we focused on a subset of mothers (n = 174) and infants (n = 166) with near-complete longitudinal sampling (Methods). Within this subset, the proportion of persistent vOTUs was significantly higher in mothers compared to infants (median = 34.3% vs. 11%, p = 4.08 × 10^−31^, Wilcoxon rank-sum test) (**Extended Data Fig. 7b**), reflecting the trend across the full cohort and consistent with a greater virome stability in adults. This difference was also evident at abundance level, as persistent vOTUs accounted for a mean 63.5% of the total virome in mothers, compared to a mean of 35.5% in infants (p = 1.4 × 10^−14^, LMM) (**Fig. 4b**). Interestingly, the proportion of persistent viruses was higher in infants delivered at home (β = 0.04, p = 0.04) (**Fig. 4c**), suggesting a potential influence of the place of delivery on virome persistence (**Supplementary Tables 26 and 27**).

Next, we investigated which phage mechanisms might contribute to their persistence in the infant and maternal gut. First, we hypothesized that a temperate viral lifestyle could help phages persist longer. After adjusting for vOTU abundance, we found that the viral lifestyle was significantly associated with persistence in mothers (FDR = 2.08 × 10^−10^, GLM) but not infants (FDR = 0.48, GLM) (**Fig. 4d**) (**Supplementary Tables 28 and 29**). We then explored whether phage ADS, which counteract bacterial defenses, could facilitate persistence. The presence of ADS was initially associated with vOTU persistence in both infants (FDR = 3.32 × 10^−3^, GLM) and mothers (FDR = 1.26 × 10^−4^, GLM) (**Supplementary Tables 28 and 29**). However, after accounting for predicted bacterial hosts as a random effect, the association was no longer significant in infants (FDR = 0.43, GLMM) (**Supplementary Table 30**). When modeling the number of infants in which each vOTU persisted, we found that anti-RM systems (FDR = 3.46 × 10^−10^) and the overall presence of ADS (FDR = 1.14 × 10^−5^, zero-inflated negative binomial GLMM) were strongly associated with longer persistence in the gut (**Supplementary Table 31**). In mothers, ADS remained significantly associated with both binary persistence (FDR = 1.25 × 10^−2^, GLMM) and the extent of persistence across individuals (FDR = 1.39 × 10⁻^7^), after controlling for host genus, with the strongest effect also driven by the presence of anti-RM systems (FDR = 1.59 × 10^−13^, zero-inflated negative binomial GLMM) (**Supplementary Tables 32 and 33**). Specifically, the DNA adenine N6-methyltransferase *hin1523* (targeting Type II RM systems) was the anti-RM subtype primarily driving the strong associations with viral persistence (**Fig. 4e**). Together, these findings indicate that long-term viral colonization is shaped not only by bacterial host but also by phage-intrinsic features, with viruses carrying ADS, particularly anti-RM systems, showing broader persistence in both infants and adults.

### DGRs are widely active and associated with phage persistence in the infant and maternal gut

To identify additional drivers of phage persistence in the developing infant gut, we investigated the role of DGRs, which have been proposed to enhance phage adaptability and persistence^32,37,62^.We identified DGRs in 2,548 vOTU representatives, including 1,004 from infants, with 39 genomes harboring multiple putative DGR systems. These DGRs targeted 2,930 distinct genes (including 336 DGRs with multiple predicted targets), the majority of which lacked functional annotation (**Supplementary Table 34**). Among the annotated targets, phage tail proteins were the most common (n = 825), followed by head and packaging proteins (n = 250) (**Fig. 5a**). Notably, we found no DGRs targeting known anti-defense proteins. While the proportion of DGR-encoding vOTUs (DGR+) detected in mothers remained stable during and after pregnancy (**Extended Data Fig. 7c**), it increased markedly in infants over the first year of life (β = 3.86 per month, p = 2.29 × 10^−20^, LMM) (**Extended Data Fig. 7c**) and was associated with delivery mode and, at nominal significance, food allergy development. Specifically, vaginally born infants (FDR = 4.08 × 10⁻^4^, LMM) and those who developed food allergies during the first year of life (p = 0.02, LMM) showed a higher proportion of DGR+ vOTUs (**Fig. 5b**) (**Supplementary Tables 35 and 36**). The presence of DGRs was significantly associated with vOTU persistence in both mothers (FDR = 1.06 × 10⁻^9^) and infants (FDR = 5 × 10^−5^, GLMM) (**Supplementary Tables 30 and 32**). Moreover, maternally shared vOTUs were significantly more likely to encode DGRs (p = 5.23 × 10⁻^15^, GLM) and showed increased persistence in infants (p = 3.08 × 10⁻^36^, GLMM) (**Extended Data Fig. 7d,e**). However, the association with DGR presence was mostly driven by *Bacteroides* phages (p = 0.61, GLMM with bacterial host as a random effect), which accounted for 51.3% of shared vOTUs and 35.7% of DGR-encoding vOTUs, suggesting a potential host-driven effect in addition to (or instead of) a potential role of DGRs in promoting their long-term maintenance in the infant gut.

**Fig. 5.**
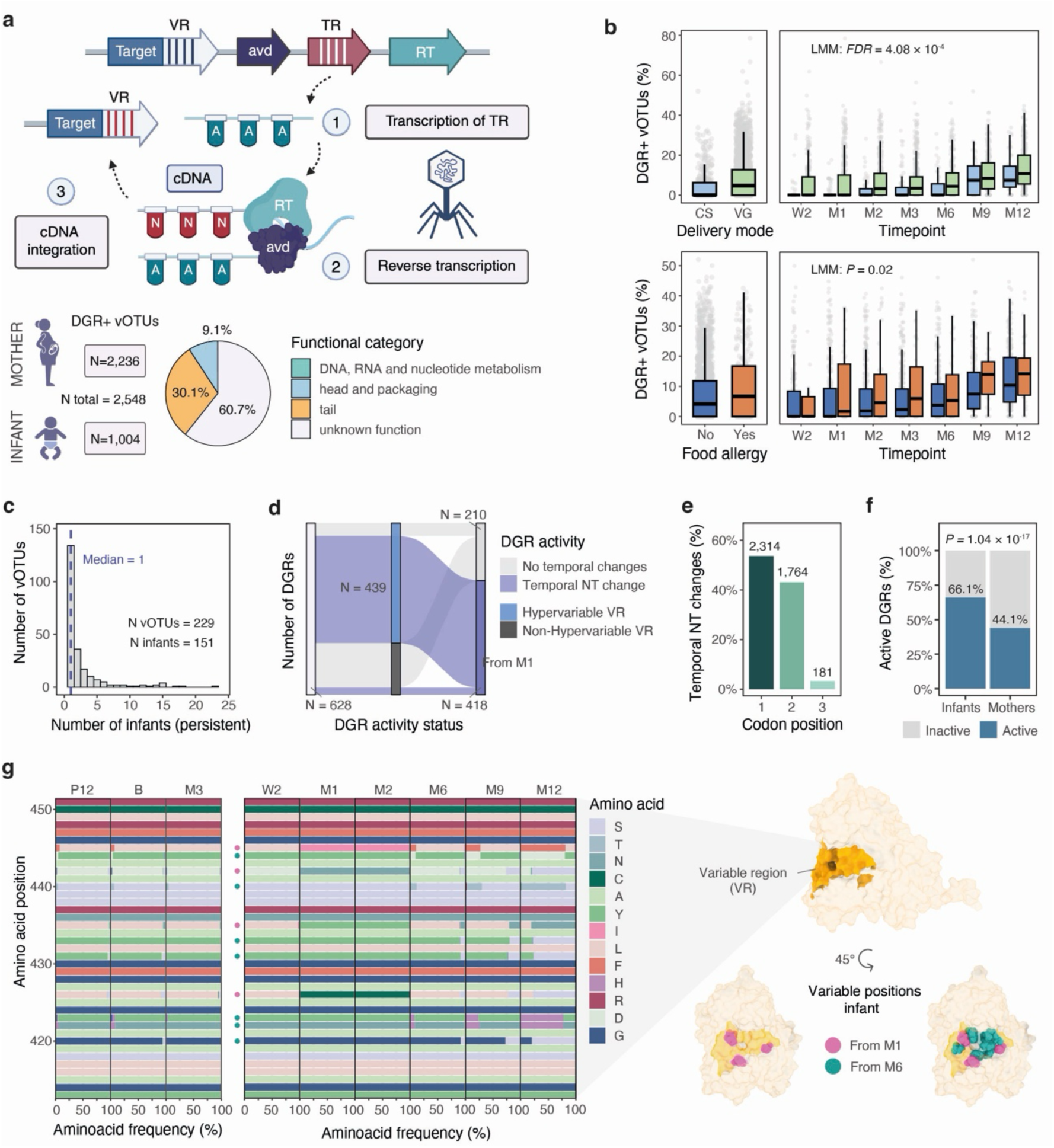
Diversity-generating retroelements are common in the gut virome and highly active in infants. **(A)** Schematic overview of the structure and mechanism of diversity-generating retroelements (DGRs). Top panel shows the genomic organization of a typical DGR system, including the target gene containing the variable region (VR) that undergoes mutagenesis, the accessory variability determinant (Avd), the template region (TR) and the error-prone reverse transcriptase (RT). The mechanism of DGR-mediated mutagenic retrohoming is shown below and includes: (1) transcription of the TR to produce TR-RNA, (2) error-prone reverse transcription by the DGR-encoded RT, characterized by adenine-specific mutagenesis, and (3) replacement of the VR alleles with the mutagenized cDNA, resulting in localized hypervariation. At the bottom of the panel, the number of DGR-encoding vOTUs identified in mothers and infants is shown, along with the functional category annotations of their target genes. **(B)** Comparison of the proportion of DGR-encoding vOTUs (DGR+) in infants according to delivery mode (top) and food allergy status (bottom). Comparisons are shown across all timepoints (left) or at individual timepoints (right). Boxplots show median, IQR and 1.5 × IQR whiskers. Gray points represent individual samples. **(C)** Histogram showing the distribution of the number of infants in which individual vOTUs persist over time. The dashed blue line indicates the median number of infants in which vOTUs persist (median = 1). **(D)** Alluvial plot summarizing DGR activity status in infants, classified according to the presence of higher nucleotide variability in VR vs non-VR and the detection of temporal nucleotide changes in the VR. **(E)** Proportion of temporal nucleotide changes occurring at each codon position across all DGR-associated VRs in infants. Absolute counts are indicated above the bars. **(F)** Comparison of the proportion of active DGRs (showing temporal nucleotide changes) in infants and mothers. **(G)** Temporal amino acid diversification within the VR of the DGR target gene from the LLNEXT_24455 vOTU in a mother–infant pair in which strain-sharing of this vOTU was detected. On the right, the predicted protein structure is shown, highlighting the VR and the corresponding variable amino acid positions. Variable positions are colored pink when diversification begins at month 1 and blue when it begins at month 6.

We next assessed DGR activity using two complementary approaches. We first quantified within-sample nucleotide variability by measuring per-site sequence diversity in the variable regions (VRs) of DGR target genes using mapped sequencing reads as a proxy for ongoing DGR-mediated diversification. In addition, the longitudinal nature of our data allowed us to explore temporal diversification by comparing VR sequences across samples collected from the same individual over time. We focused on the subset of 229 DGR+ vOTUs classified as persistent across 151 infants, representing a total of 628 DGRs (717 VRs) (**Fig. 5c**). Within this set, we identified 439 potentially active DGR systems (75.4%) (475 VRs) based on elevated nucleotide diversity at VR positions compared to non-VR regions of the target genes (FDR < 0.05, Wilcoxon test), reflecting a high level of DGR activity in the infant gut virome, consistent with previous observations^63^ (**Supplementary Table 37**). Analysis of temporal changes in consensus VR sequences showed that 392 of the potentially active DGR systems (89.5%) showed temporal nucleotide diversification in the VRs (418 in total) (**Fig. 5d**). Notably, mutations at the VR in infant phages occurred predominantly at the first and second positions of codons (β = 3.86, p = 1.29 × 10⁻^89^, GLMM), resulting in a strong enrichment of non-synonymous changes (**Fig. 5e**). Viral features and infant phenotypes were not associated to DGR activity, including phage lifestyle (virulent vs temperate: β = 0.454, p = 0.11), infant delivery mode (vaginal vs CS: β = −0.366, p = 0.45) or exclusive breastfeeding duration (β = 0.59, p = 0.12, GLMM) (**Supplementary Table 38**). To investigate whether the high DGR activity patterns were exclusive to infants, we performed the same analysis in mothers (**Extended Data Fig. 7f**). Across 1,076 persistent DGR+ vOTUs in 448 mothers (3,825 VRs), we observed a substantially lower proportion of DGRs showing temporal nucleotide changes in the VRs relative to infants (β = −1.08, p = 1.04 × 10^−17^, GLMM) (**Fig. 5f**), indicating lower DGR-mediated diversification in the adult gut virome (**Supplementary Table 39**).

Finally, we evaluated whether maternal DGRs contribute to phage adaptation in the developing infant gut. For this, we focused on 99 maternally shared DGR+ vOTUs (201 total DGRs). Shared vOTUs showed only modest but non-significant increases in the proportion of DGRs exhibiting longitudinal amino acid changes compared to non-shared vOTUs (β = 0.44, p = 0.13, GLMM) (**Supplementary Table 40**). Within this shared subset, the probability of DGR-mediated amino acid diversification was significantly higher in infants than in mothers (β = 0.85, p = 4.97 × 10^−5^, GLMM) (**Supplementary Table 41**). Consistent with this, when examining temporal DGR-driven mutations within mother–infant pairs (across 93 shared DGRs (102 VRs) detected in 45 pairs), we found that a notable proportion only started generating amino acid changes after transmission to the infant (21.6% vs. 11.8% with longitudinal amino acid changes exclusively in mothers). For example, in one mother-infant pair, a protein from a prevalent predicted *Bacteroides* phage (LLNEXT_24455, detected in 104 mothers and 56 infants) with homology to a tail fiber protein and targeted by a DGR system, remained largely unaltered during pregnancy in the maternal gut but rapidly accumulated novel amino acid variants across multiple VR positions in the infant gut. These amino acid changes were concentrated at exposed residues of the predicted structure, consistent with selection for altered receptor-binding specificity (**Fig. 5g**). Overall, these results reveal that DGRs are widespread in maternal and infant gut phages and show higher activity in infants, rapidly diversifying VR sequences and playing a key role in driving phage adaptability and persistence during early gut colonization.

## Discussion

Early life represents a critical window for microbial ecosystem assembly, yet the origins, dynamics and persistence mechanisms of the developing gut virome remain poorly defined. Through longitudinal metagenomic profiling of 714 mother–infant pairs during pregnancy and the first year of life, we generated an extensive reference catalog of maternal and early-life gut viruses. We further identified known and previously undescribed factors that substantially influence viral composition and temporal trajectories in infants and characterized multiple maternal sources of infant gut viruses. Finally, we provide evidence of viral genetic mechanisms that enhance their ability to adapt and persist within the gut environment.

Despite an increasing number of early-life viral studies^19,22,23,26,27,32,64^ and efforts to create standardized and comprehensive viral databases^37,65,66^, the immense diversity and high individual specificity of the gut virome make it difficult to rely exclusively on reference genomes for viral profiling. By leveraging our extensive gut and breastmilk metagenomic dataset and applying an assembly-based viral discovery approach, we identified 16,540 vOTUs that do not cluster with any of the reference genomes included in the study, with the >8,500 vOTUs achieving at least 90% genome completeness potentially representing novel viral species according to MIUVIG standards^17^. Characterization and profiling of the gut viral catalog in mothers and infants from our cohort highlighted the stability of the maternal virome, with no viral groups showing significant changes in abundance during pregnancy, and only minor differences observed when including post-pregnancy timepoints. While few studies have investigated the maternal pregnancy virome^22^, this stability may reflect the nature of our generally healthy cohort, which may have limited our ability to detect potential differences that could appear in more heterogeneous populations or high-risk pregnancies. This stability contrasts with the rapid viral diversification we observed in infants during the first year of life, consistent with previous reports based on smaller sample sizes^22,27,32^.

Factors shaping the developing gut virome have so far been understudied, with most previous work focusing on delivery mode and infant feeding practices^22,23,27,67^. Consistent with these studies, we found that both factors are major drivers of viral compositional trajectories during the first year of life, with strong influences on viral diversity and composition. CS delivery was associated with a strong reduction in predicted *Bacteroides* and *Parabacteroides* phage abundance, reflecting the depletion of these bacterial taxa in CS-born infants^40^, and an enrichment of phages infecting potentially pathogenic genera such as *Enterococcus*, *Enterobacter*, *Klebsiella* and *Flavonifractor*. Notably, infants who were never breastfed showed increased abundance of *Akkermansia* phages, in line with previous observations of *Akkermansia* enrichment in formula-fed infants^68,69^. Beyond these well-established influences, the inclusion of an extensive set of maternal and infant phenotypes revealed previously undescribed modulators of virome development, including maternal parity, infant birth weight, gestational age, maternal education level and infections during pregnancy. Interestingly, infants who developed food allergies during their first year showed increased viral diversity that was not fully explained by bacterial diversity alone, suggesting virus-specific contributions that require further investigation to better understand the underlying mechanisms.

Only a few studies have investigated the origin of the infant gut virome^22,24,27,32^, and these have been often limited by insufficient maternal sampling and small cohort sizes. Consistent with previous observations, we confirm that the maternal gut constitutes a major source of infant gut viruses, with mother–infant pairs displaying highly similar viral strain profiles indicating extensive strain-sharing. This strong similarity enabled highly accurate prediction of mother–infant relatedness based solely on strain profiles.

Given our longitudinal strain-level comparisons across nearly 2,000 vOTUs, we revealed that the viral-sharing rate decreased over time during the first year of life, likely reflecting the growing influence of non-maternal environmental exposures, and that viral sharing occurs less frequently in CS-delivered pairs. Ultra-deep sequencing of breastmilk metagenomes further uncovered this niche as an additional maternal source of infant gut viruses, albeit one less dominant than the maternal gut, indicating that multiple maternal reservoirs contribute to early virome establishment. Future efforts to generate extensive, deep metagenomic profiles of this and other understudied maternal niches such as the vaginal and oral microbiomes could reveal their full contribution to the infant gut virome.

Supporting the hypothesis of viral transmission via co-transferred bacterial hosts^22,24,27^, we show that temperate phage and bacterial host co-transmission is common, accounting for approximately 15–20% of temperate phage-sharing events. This proportion is likely an underestimate given challenges in recovering low-abundance MAGs from metagenomic reads and variable MAG completeness. Finally, through combined sequence- and structure-based analyses, we identified numerous viral PFs enriched among shared viruses, including structural proteins and others with accessory or auxiliary functions, such as prevalent ADS in infants (e.g., anti-Thoeris) and additional PFs recently reported to have defense or anti-defense roles^58–61^. Further experimental validation is needed to confirm the functional roles of shared viral proteins.

While initial viral seeding is critical for establishing the early virome and influencing early host–microbe interactions, the long-term impact of viruses on the gut ecosystem depends not only on their origin but also on their ability to persist. However, the ecological and genetic factors that enable phages to establish stable, long-term associations with their hosts remain poorly understood, particularly in the highly dynamic environment of early life. By examining the relationship between viral genetic features and persistence, we found that a temperate lifestyle, reflecting the ability of viruses to reside as part of dormant (lysogenic) infections into bacterial cells, was associated with persistence in the maternal but not the infant gut. This difference likely reflects the distinct ecological context of early life, characterized by rapid bacterial succession and ongoing microbial colonization^70,71^, which may limit the long-term advantages of lysogeny. In contrast, viral mechanisms that facilitate rapid adaptation were strongly associated with long-term viral maintenance in both mothers and infants. In particular, adenine methyltransferases, which methylate phage DNA to evade host Type II restriction modification systems^72^, and DGRs, which generate diversity in phage receptor-binding proteins through targeted, error-prone reverse transcription^73^, were highly prevalent and strongly linked to viral persistence. Both methyltransferases and DGRs have recently been proposed as key determinants enabling broader viral host range^37^. Notably, the proportion of DGR-encoding phages was higher among vaginally born infants and those who developed food allergies during the first year of life. Furthermore, we found that DGRs were highly active in the gut, showing significantly higher temporal nucleotide diversification in infants than in mothers, indicative of rapid phage adaptation and intensified phage–host co-evolution during early life. In agreement with prior work^34^, these findings suggest that gut phage persistence, particularly during early development, is driven less by stable genomic integration and more by continual phage–host co-evolution, consistent with a Red Queen–type arms race^74^.

Despite extensive characterization of the maternal and infant gut virome, our study is mostly restricted to double-stranded DNA viruses, as our methods did not capture the active virome fraction via virus-like particle isolation, so single-stranded DNA and RNA viruses were not assessed. Additionally, the relative homogeneity of the Lifelines NEXT cohort may have limited our ability to detect or fully capture the strength of certain phenotype associations, such as the subtle associations we observed between viral diversity and infant food allergy. Validation through meta-analyses or in more diverse cohorts will be important to confirm our findings and expand our understanding of early-life virome–host interactions.

Together, our findings provide a comprehensive view of pregnancy and early gut virome dynamics, highlighting the maternal contribution and viral genetic mechanisms as key factors shaping early-life gut viral assembly and long-term colonization.

## Methods

### Study population and metadata collection

Samples were obtained from the LLNEXT cohort^38^, a prospective birth cohort designed to investigate intrinsic and extrinsic determinants of early-life health and development, nested within the Lifelines population-based cohort in the Northern Netherlands^75^. LLNEXT recruited pregnant women, their partners and infants between 2016 and 2023 for longitudinal collection of biological samples, including feces and breastmilk, across 10 timepoints, alongside extensive clinical and lifestyle metadata. A detailed description of the study design and collected predictors is provided in our previous publications^7,22^. The present study included the first 714 mother–infant pairs recruited between 2016 and 2019, comprising 696 first pregnancies, 18 second pregnancies and 12 sets of twins. Of the available metadata, 42 phenotypes capturing clinical, perinatal, dietary, socioeconomic, environmental, and infant health and growth–related information were included in our downstream analyses (**Table S1**). The study was approved by the UMCG Ethics Committee (METc2015/600), and all participants provided written informed consent.

### Sample collection

#### Fecal samples

The maternal (n = 1,587) and infant (n = 2,936) fecal samples in the present study were collected as described in our previous publication^7^. In short, mothers collected their own samples during pregnancy (P12 and P28), delivery (B) and 3 months postpartum (M3). Parents or guardians collected infant fecal samples starting from the second week of life (W2) up to 12 months (M12), with an average of four samples and maximum of seven samples collected per infant (Figure 1A). All samples were collected using stool collection kits provided by the UMCG and frozen at home at −20°C within 10 min of stool production. Frozen samples were then collected by UMCG personnel, transported under frozen conditions and subsequently stored at −20 °C for short-term or −80 °C for long-term storage until DNA extraction.

#### Milk samples

Human milk samples were collected as previously described^45^. For this study, 91 samples collected at M1 were included. Briefly, mothers were asked to pump milk from one breast using their usual breast pump, collecting milk from the second feeding after midnight and at least 2 hours after the previous feeding. After gentle shaking to ensure homogenization, 2 ml aliquots were prepared in cryotubes (122279, Greiner Bio-One, Kremsmünster, Austria) using plastic dropper pipettes (H10041, MLS, Menen, Belgium). Samples were stored at −20 °C at home for up to 6 months, transported to UMCG under frozen conditions and stored at −20 °C for short-term or −80 °C for long-term storage until processing.

### DNA extraction

Fecal microbial DNA was extracted from 0.2–0.5 g of fecal material using the QIAamp Fast DNA Stool Mini Kit (Qiagen, Germany) and the QIAcube automated system (Qiagen), following the manufacturer’s instructions. Extractions were performed at the Institute for Clinical Molecular Biology (Kiel, Germany) with a final elution volume of 100 μl. DNA eluates were subsequently stored at −20 °C. Milk microbial DNA was extracted from 3.5 ml of milk using the DNeasy PowerSoil Pro Kit (47016, Qiagen, Venlo, Netherlands), as previously described^45^. Briefly, milk samples were centrifuged at 13,000 × g at 4° C for 15 min, removing fat and supernatant. Pellets were resuspended in 800 µl Solution CD1, transferred to PowerBead Pro Tubes and vortexed briefly. Samples were incubated at 65° C for 10 min, followed by bead beating at 5000 rpm at 4° C for 45 s using a Precellys Evolution Homogenizer (Bertin Instruments, Rockville, Maryland, USA). The lysates were centrifuged again at 15,000 × g at 4° C for 1 min, and DNA was extracted from the supernatant using the *DNeasy PowerSoil Pro Kit with Inhibitor Removal Technology Protocol* (version May 2018) on a QIAcube (Qiagen) with a final elution volume of 50 µl. DNA extracts were stored at −20° C.

### Library preparation and sequencing

Fecal and breastmilk microbial DNA was sent to Novogene (Cambridge, UK) for library preparation and shotgun metagenomic sequencing. Libraries were prepared using either the NEBNext® Ultra™ DNA Library Prep Kit or the NEBNext® Ultra™ II DNA Library Prep Kit, depending on DNA concentration. Sequencing was performed on the Illumina HiSeq 2000 or NovaSeq 6000 platforms using 2 × 150 bp paired-end chemistry.

### Sequencing data processing and *de novo* assembly

Shotgun metagenomic reads were processed to remove low-quality sequences and human DNA contamination. Adapters and low-quality bases were trimmed using BBDuk (v39.01) with ‘ktrim=r k=23 mink=11 hdist=1 tpe tbo’ parameters and KneadData (v0.10.0) with ‘--trimmomatic-options “LEADING:20 TRAILING:20 SLIDINGWINDOW:4:20 MINLEN:50” --bypass-trf --reorder’ parameters and default quality-scores. Human reads were filtered against the GRCh38 reference genome. The quality of processed reads was assessed with FastQC (v0.11.9). Quality-filtered reads from each sample, including unmatched reads, were assembled into contigs using MetaSPAdes (v3.15.5) with default parameters^76^.

### Viral identification workflow

Metagenome-assembled contigs >10 kb were screened for viral origin using VirSorter2^77^ (v2.2.4), DeepVirFinder^78^ (v1.0) and geNomad^79^ (v1.5.1). Contigs were retained if they met at least one of the following thresholds: VirSorter2 viral score > 0.5, DeepVirFinder viral score ≥ 0.95 with p < 0.01 or classification as viral by geNomad. VirSorter2 was run with the --keep-original-seq and --include-groups “dsDNAphage,NCLDV,ssDNA,lavidaviridae” options, DeepVirFinder with default settings and geNomad with --enable-score-calibration and --disable-find-proviruses. Potential viral contigs were combined and processed with COBRA^80^ (v1.2.2) to generate assemblies with improved contiguity and completeness. The default end-to-end geNomad pipeline was then applied to ensure viral origin and trim potential flanking host regions of extended sequences.

Next, CheckV^81^ (v1.0.1) was used to: (i) further remove residual host contamination with the “contamination” module and (ii) assess genome quality with its end-to-end workflow. Only contigs meeting the following thresholds were retained: ≥ 50% estimated completeness, viral-to-host gene ratio >1 and a total genome length less than 1.5 × the expected size. Predicted viral sequences were then combined with viral genomes from multiple public databases: MGV^65^ (n = 189,680 genomes with >50% estimated completeness), GPD^66^ (n = 82,621 genomes with >50% estimated completeness),^76^ IMG/VR^82^ (n = 36,064 genomes with >50% estimated completeness from the *Duplodnaviria* realm identified in human-associated samples),^77^ ELGV^64^ (n = 21,295 genomes with >50% estimated completeness),^21^ Viral RefSeq^83^ (n = 5,199 genomes >10 kb after excluding the *Riboviria* realm), Shah et al.^84^ (n = 4,627 dsDNA viruses with >50% estimated completeness), Benler et al.^85^ (n = 1,480 viral genomes belonging to the *Uroviricota* phylum)^79^ and CrAss-like phage databases (genomes with >50% estimated completeness from Gulyaeva et al.^86^ (n = 637) and Yutin et al.^87^ (n = 648) and all viral genomes from Guerin et al.^88^ (n = 249)). All sequences were then clustered at 99% ANI and 85% alignment fraction (AF) using the CheckV scripts *anicalc.py* and *aniclust.py* with the ‘--min_ani 99 --min_tcov 85 --min_qcov 0’ parameters. Any clusters containing sequences detected in negative control samples (n = 2) were removed from further analysis. The remaining viral genomes were dereplicated using the MIUVIG-recommended thresholds^17^ (95% ANI, 85% AF). For each cluster, the longest sequence was selected as the vOTU representative. Maternal breastmilk samples were processed in the same manner, with LLNEXT-GV representative genomes included in the dereplication step. Genus- and family-level vOTUs were defined by clustering the representative viral genomes using a combination of pairwise average amino acid identity and gene-sharing profiles, following the approach of Nayfach et al^65^. In addition, viral sequences generated in this study were further deduplicated (99% ANI, 95% AF) and dereplicated (95% ANI, 85% AF) separately to generate a non-redundant set of viral genomes and representative sequences derived exclusively from this dataset, independent of external databases.

### Viral abundance estimation

To estimate the viral abundances in each sample, we mapped the metagenomic reads, filtered and quality-trimmed as described above, to the database of vOTU-representative sequences generated using Bowtie2^89^ (v2.4.5) with --very-sensitive --no-unal’ options. Breadth of genome coverage by sequencing reads was calculated using the BEDTools^90^ (v2.30.0) command *coverage*. Each vOTU was considered present in a sample if the mapped reads covered >75% of the genome length, with their relative abundances expressed in reads per kilobase per million mapped reads (RPKM). Representative vOTUs detected as present in fecal and breastmilk metagenomes were used to define the LLNEXT-GV and LLNEXT-BMV catalogs, respectively.

### Taxonomy assignment

Taxonomic classification of vOTUs was performed using a hierarchical approach. First, vOTUs containing viral RefSeq genomes were directly assigned taxonomy based on RefSeq annotations. When multiple RefSeq genomes with differing taxonomy were clustered within the same vOTU, the lowest common taxonomic rank was assigned. For vOTUs lacking RefSeq matches, taxonomy was inferred using geNomad (v1.11.0) and VITAP^91^ (v1.7.1), which align viral proteins against curated marker sets and reference databases encompassing ICTV-recognized lineages^92^. When only one tool produced an assignment, that taxonomy was retained. When both were available, we used VITAP by default unless the two annotations were consistent up to the class level and geNomad provided a more complete taxonomy, in which case the geNomad assignment was selected.

### Prediction of bacteriophage lifestyle

The lifestyle of bacteriophages (temperate or virulent) was inferred using BACPHLIP^93^ (v0.9.6), which classifies phages based on the presence of lysogeny-associated protein domains. Given the variability in phage genome completeness, lifestyle assignments were made following previously established criteria^94^. Phages with BACPHLIP scores >0.5 were classified as “temperate”. “Virulent” classification was only applied to phages with scores >0.5 and ≥ 90% estimated genome completeness (as determined by CheckV), to avoid misclassification due to potential undetected lysogeny-associated domains in incomplete genomes. Eukaryotic viral genomes were excluded from lifestyle prediction.

### Viral host prediction

Putative bacterial hosts were assigned to vOTUs at genus level using the iPHoP framework^95^ (v1.3.3). Host prediction was performed using both the default reference database (iPHoP_db_Aug23_rw) and a custom-expanded database that incorporated dereplicated MAGs generated in this study. For each vOTU, the host genus with the highest confidence score across both databases was retained, applying a minimum prediction confidence threshold of 90 (default). When multiple host predictions shared the highest confidence score, the prediction corresponding to their lowest common ancestor was selected.

### Identification and analysis of viral ADS

To characterize viral ADS, we analyzed all vOTU-representative genomes using DefenseFinder^52^ (v2.0.0) with the DefenseFinder models database v2.0.2. The analysis was performed with the --db-type gembase option and all other parameters set to default. Enrichment of specific ADS in maternal versus infant vOTUs was assessed using Fisher’s exact tests, including only systems with at least 20 occurrences across all individuals. Enrichment analyses of anti-RecBCD and anti-CBASS by predicted bacterial hosts were performed using the same approach, including only those genera with vOTUs carrying at least 50 identified systems.

### Bacterial abundance profiling and binning

Bacterial abundance profiling and binning in LLNEXT gut metagenomic samples were performed as previously described^7^. Briefly, taxonomic composition was estimated using MetaPhlAn4^96^ with the MetaPhlAn database of marker genes mpa_vJan21 and the ChocoPhlAn species-level genomic bin (SGB) database (202103). Bacterial binning was performed separately for each metagenome using metaWRAP^97^ (v1.3.2), and MAGs were retained if they had a minimum completeness of 50% and a maximum contamination of 10%, as estimated by CheckM^98^ (v1.0.12).

### Viral diversity and composition in mothers and infants

Viral richness and alpha diversity (represented by the Shannon diversity index) of each metagenomic sample was estimated using the specnumber and diversity functions of the vegan package^99^ (v2.7-1). The acquisition of new vOTUs per infant sample was defined as the number of vOTUs detected in that sample but absent from all previously collected samples from the same individual. Associations of viral metrics with time (modeled as a continuous variable in months) were performed using LMM with the lmerTest package^100^ (v3.1-3), including read depth and DNA concentration as covariates and individual ID as a random effect. Beta diversity was calculated using Bray-Curtis distances based on vOTU abundances, and temporal shifts in virome composition were evaluated using PERMANOVA with 1,000 permutations, restricting permutations within individuals to account for repeated measurements (‘strata’ parameter), including read depth and DNA concentration as covariates, and applying “by = margin” (adonis2 function, vegan package). Non-metric multidimensional scaling (NMDS) was additionally performed to visualize patterns of viral community composition. To compare the virome similarity (calculated as 1 minus Bray-Curtis dissimilarity) between (1) within-mother and within-infant samples and (2) across levels of biological-relatedness, we performed a non-parametric permutation test (10,000 permutations) in which group labels were randomly permuted. Associations between viral abundances and time were assessed using LMM, as described above, for both vOTUs and viral groups (by predicted bacterial host genus) after log transformation, adding half of the minimum overall abundance value as a pseudocount. For this analysis, time was encoded as a categorical variable representing each sampled timepoint, using W2 and P12 as references for infants and mothers, respectively. Only vOTUs and viral groups present in ≥ 5% of infant samples or ≥ 10% of maternal samples were included in the association analysis. Significance of the associations was assessed using likelihood ratio tests comparing full models (including time) to null models (without time). Overall viral prevalence in mothers and infants was assessed using a paired Wilcoxon test comparing the number of maternal and infant vOTUs above each prevalence threshold.

### Association of infant viral features with human phenotypes

Association between human phenotypes and infant viral features (Shannon diversity, richness, relative abundance of temperate phages and log-transformed, prevalence-filtered viral abundances) were assessed using LMM, as described above, additionally including timepoint as a covariate. For viral abundances grouped by predicted bacterial host, models were further adjusted for the corresponding bacterial host abundance at the genus level (CLR-transformed), converted from Metaphlan SGB-based taxonomy to GTDB using the sgb_to_gtdb_profile.py script. Associations with viral composition based on viral abundances grouped by host were assessed using PERMANOVA (10,000 permutations) per timepoint, as previously described (excluding the ‘strata’ parameter).

To explore temporal trajectories of the gut virome, we applied TEMPTED^41^, a time-informed dimensionality reduction method for longitudinal microbiome studies. This analysis was restricted to infants with samples collected at three or more time points (n = 473). Viral abundances grouped by predicted host were log-transformed and used as input, and TEMPTED was run with ‘transform = “none”, r = 5’, keeping all other parameters at default. Inferred subject trajectories (PC1‒PC5) were then associated to human phenotypes using LMM. Given the known effects of delivery and feeding mode on the infant gut microbiome, all associations (including LMM and PERMANOVA) were performed both without and with adjustment for these biological covariates.

### Viral strain-sharing analysis

Viral vOTU representatives detected in at least one fecal or breastmilk metagenome (n = 31,205) were remapped to the metagenomic reads, and sequence alignment files were processed using the inStrain^101^ (v1.9.0) profile function, with a minimum mapQ score of 0 and insert size set to 160. Genome similarity among samples with a vOTU genome breadth ≥ 75% was assessed using the inStrain compare function. Only regions with ≥ 5x coverage were included in comparisons, and sample pairs with < 75% comparable regions across phage genomes were excluded. Population-level ANI (popANI) values were used to assess strain-similarity within each vOTU, considering phage strains as shared between samples if their genomic regions exhibited > 99.999% popANI. Strain-similarity in related versus unrelated mother–infant pairs was assessed using a one-sided permutation test with 10,000 permutations. The proportion of strain-sharing events was compared using chi-squared tests. To evaluate whether viral strain-similarity profiles could accurately identify biological mother–infant pairs, logistic regression models were trained using popANI-based similarity values between maternal samples (any timepoint) and each infant timepoint. All models (one per infant timepoint) were implemented within the tidymodels framework^102^ in R (v1.3.0), using logistic regression fitted with the glm engine. Model performance was evaluated using 10 repeats of 5-fold cross-validation, where each fold used 80% of the data for training and 20% for testing. Predictions from all folds and repeats were aggregated to construct a single receiver operating characteristic (ROC) curve, and the resulting AUC was computed with the pROC package^103^ (v1.19.0.1). Longitudinal analyses of the proportion of families with detected strain-sharing and the strain-sharing rate, defined as the fraction of infant vOTUs shared with the corresponding mother, were performed using GLMMs with family ID as a random effect.

### Comparison of breastmilk and fecal sample diversity and strain-sharing

The viral diversity in breastmilk (M1) was compared to the closest maternal fecal samples (B) from the same mothers (n = 91) using a paired Wilcoxon test. Comparison of viral strain-sharing via breastmilk versus maternal feces to the infant gut was performed using McNemar’s test. Due to the low number of breastmilk–infant sharing events (n = 12), temporal trends in strain-sharing were not statistically analyzed.

### Temperate phage‒bacterial host co-sharing analysis

The association between temperate viral lifestyle and maternal–infant viral strain-sharing was assessed using a GLM with a binomial distribution, including maternal mean abundance of each vOTU as a covariate. To investigate potential co-transmission, all potentially shared temperate vOTUs (n = 435, popANI > 99.999% in at least one mother–infant pair) were mapped to maternal and infant MAGs from samples of individuals in whom strain-sharing of these vOTUs was detected (n = 26,106 MAGs with at least medium-quality, completeness ≥ 50% and contamination ≤ 10%). Mapping was performed using minimap2^104^ (v2.26) with parameters “-x asm5”, retaining hits with ≥ 75% genome coverage and ≥ 95% sequence identity. For each vOTU, mapping results were compared with predicted bacterial host assignments at the genus level to assess concordance. Potential maternal–infant co-sharing events were defined as cases where the same temperate vOTU mapped to MAGs from both members of a mother–infant pair. The enrichment of detected co-sharing events was tested using a permutation test against 10,000 randomly permuted mother–infant pairs. Confirmation of strain-level co-sharing was performed by comparing ANI between paired maternal and infant MAGs using SKANI^105^ (v0.3.0) with triangle command, defining strain-level sharing as ANI > 99.9%.

### Clustering of viral proteins into PFs

Open reading frames (ORFs) were predicted from all representative viral sequences obtained from fecal and milk metagenomes (n = 1,811,012) using Prodigal-gv^106^ (v2.11.0). The resulting protein sequences were clustered into non-redundant PFs (n = 282,228) using MMseqs2^107^ (v15.6f452) through a two-step clustering strategy adapted from Camargo et al.^37^. First, proteins were clustered based on at least 80% bidirectional coverage using mmseqs cluster with the parameters: “-s 7.0 -c 0.8 -e 1e-3 --cluster-mode 0 --cluster-reassign 1 --kmer-per-seq 50”. For each cluster, MSAs were generated using a star-center alignment (results2msa --msa-format-mode 2), from which profiles and consensus sequences were derived using msa2profile with “--msa-type 2 --match-mode 1 --match-ratio 0.5” parameters and profile2consensus. Consensus sequences were then compared to cluster profiles using a profile-to-consensus search with mmseqs search (--cov-mode 0 -c 0.9 -s 8.0 -e 1e-4 --add-self-matches 1 -a 1), a more sensitive approach than the initial protein-to-protein search. Profiles were subsequently clustered to merge related PFs into larger, more diverse clusters (mmseqs clust), and the results of both clustering steps were combined using mmseqs mergeclusters to produce the final set of PFs. MSAs for these secondary clusters were generated using the same star-center alignment method (result2msa -- msa-format-mode 2).

### Functional enrichment of PFs

To identify PFs potentially enriched in early infant viruses of maternal origin, viral proteins were first classified as shared if they were encoded by a vOTU shared at the strain level (popANI > 0.99999) between at least one maternal–infant sample pair, limited to infant samples from W2 to M3 (n_vOTUs_ = 449). Proteins from vOTUs detected in maternal samples but not shared at the strain level were classified as non-shared (n_vOTUs_ = 14,449). Only PFs containing at least 10 proteins across the tested vOTUs (shared + non-shared) were included in enrichment analysis. Enrichment was assessed using Fisher’s exact test (fisher.test R function), and PFs with a FDR < 0.05 and OR >1 were considered significantly enriched among maternally shared viruses.

### Functional annotation of enriched families

Representative sequences of enriched PFs (n = 986) were first compared against the HMM profiles of the PHROGs^50^, KOfam^51^ and AntiDefenseFinder^52^ databases. For PHROGs, HMMER^108^ hmmsearch (v3.4) was used, retaining only the most significant hit per query with an e-value < 1×10⁻^5^. For KOfam, KoFamScan^51^ (v1.3.0) was applied using default parameters, and hits were considered significant if the alignment score was equal or higher than the predefined KO thresholds. In the case of AntiDefenseFinder, DefenseFinder^52^ (v.2.0.0) was used with “--db-type gembase --antidefensefinder-only” parameters. MSAs of PFs that remained unannotated were subsequently enriched with UniRef30 sequences^109^ (release 2023_02) using HHblits^110^ (v3.3.0) with parameters “-id 80 -n 3 -e 0.1”.

Protein structures for each PF were then predicted using ColabFold^53^ (v1.5.5) with the options “--num-recycle=3 --recycle-early-stop-tolerance 0.5 --num-seeds 3”. For each PF, the best-ranking structural model according to the pLDDT score was selected, retaining only models with an average pLDDT ≥ 70. Predicted structures were then queried against the PDB^54^, AFDB^55^ and BFVD^56^ using Foldseek^57^ (v10.941cd33) in easy-search mode with the TM-align mode. Matches were considered significant if they met the following thresholds: qtm-score > 0.7, prob ≥ 0.9 and qcov ≥ 0.7. For each query, only the hit with the highest probability (prob) was retained per database. In cases where multiple hits had the same probability, the one with the highest qtm-score was selected. The top hits were subsequently annotated by retrieving corresponding UniProt information, including protein name, functional description, Gene Ontology terms, keywords and domain/family annotations. For PDB entries that mapped to multiple UniProt accessions, a single representative UniProt entry was selected based on manual curation.

### Analysis of maternal and infant phage persistence

Viral persistence was defined separately for mothers and infants based on longitudinal detection. In infants, persistent vOTUs were defined as those detected in at least three timepoints from the same individual, including at least one early (W2, M1, M2 or M3) and one late (M6, M9 or M12) timepoint. In mothers, persistent vOTUs were defined as those detected in at least one pregnancy timepoint (P12, P28 or B) and the post-pregnancy timepoint (M3) from the same individual. Comparison of the proportion and abundance of persistent vOTUs between mothers and infants was performed using a Wilcoxon rank-sum test (proportion) and a LMM with individual ID as a random effect (abundance), restricted to individuals with near-complete longitudinal sampling (excluding the B timepoint for mothers and the W2 timepoint for infants due to lower sample availability). The influence of a subset of 10 phenotypes on the proportion of persistent vOTUs was assessed using a linear model including read depth and DNA concentration as covariates. This analysis was performed with and without adjustment for feeding mode (ever/never breastfed) and delivery mode.

Next, we explored associations between viral genomic features and persistence. First, the association between temperate lifestyle, the presence of phage ADS and DGRs with binary persistence status (persistent in at least one individual or not) was assessed using GLMs adjusted for each vOTU’s mean abundance. To account for potential host-specific effects, the associations with ADS (including the five most common anti-defense system types and anti-restriction modification subtypes with at least 50 occurrences across all vOTUs) and DGRs were further assessed using binomial GLMMs with the glmmTBB package^111^ (v1.1.12), including the predicted host genus as a random effect. To test the association with the extent of persistence across individuals (number of individuals in which each vOTU persisted), zero-inflated negative binomial GLMMs were used. All GLMMs were also adjusted for the mean abundance of each vOTU.

### DGR identification and characterization

Detection of DGRs in vOTU-representative genomes was performed in three steps, following a previously established approach^62^. First, ORFs were predicted using Prodigal-gv^106^ (v2.11.0), and candidate reverse transcriptase (RT) genes were identified via HMM profile searches using hmmsearch (v3.4). Next, nucleotide repeats flanking the putative RTs were detected using BLASTn^112^ (v2.15.0) with the parameters “-word_size 8 -dust no -gapopen 6 -gapextend 2”. Finally, candidate DGRs were selected based on repeat architecture and RT domain length. In cases where multiple RTs within a vOTU were predicted to target the same VR region, only the RT closest to the target gene was retained. Systems showing adenine mutation frequencies < 75% were excluded from downstream analyses. DGR targets were annotated using HMMER^110^ hmmsearch (v3.4) against PHROGs HMM profiles. The association between the proportion of DGR-encoding vOTUs with time, as well as with a subset of 10 phenotypes, was performed using a LMM, as previously described. The association of maternal–infant viral-sharing with DGR presence was first assessed using a binomial GLM, adjusting for mean maternal vOTU abundance. This association, together with the association of sharing with viral persistence, was also analyzed using a binomial GLMM, including mean maternal (for DGR presence) or infant (for persistence) vOTU abundance as a covariate and bacterial host as a random effect.

### DGR activity analysis

DGR activity was assessed using anvi’o^113^ (v8). Briefly, a contigs database was first generated from all vOTU genomes detected in at least one fecal or breastmilk metagenome using anvi-gen-contigs-database, providing an external gene calls file derived from Prodigal-gv predictions. Individual BAM files from read mapping were then used to compute sample-specific single nucleotide variant profiles via anvi-profile with codon frequency characterization enabled (--profile-SCVs). By default, only nucleotide positions with a minimum coverage of 10X were included in the analysis. These individual profiles were merged into a single profile database using anvi-merge. To quantify variability in DGR target genes, we used anvi-gen-variability-profile at both nucleotide (--engine NT) and amino acid (--engine AA) levels. For the nucleotide level, we applied the --quince-mode option, whereas amino acid–level analyses were performed using the --kiefl-mode option. For each individual, we retained only DGR-containing vOTUs that were persistent over time, and variability profiles were computed across that individual’s longitudinal samples. DGR activity was determined by comparing the per-site nucleotide diversity (entropy) in the VR versus non-VR region within each sample using a one-sided Wilcoxon rank-sum test and by assessing the occurrence of nucleotide or amino acid substitutions at the same genomic positions in the VR across multiple time points from the same individual. Position bias of DGR mutations was assessed using a binomial GLMM, with both vOTU ID and individual ID included as random effects, comparing the frequency of substitutions at the 1^st^ and 2^nd^ codon positions versus the 3^rd^. GLMMs were also used to evaluate associations between DGR activity (presence of longitudinal nucleotide changes) and viral (lifestyle) or infant (delivery and feeding mode) features and to assess DGR-mediated nucleotide and amino acid diversification across various comparisons. The protein structure of the DGR target gene in the LLNEXT_24455 vOTU was predicted using AlphaFold2^114^ (v2.3.1), based on the infant consensus protein sequence and using the monomer preset. Searches were performed against the default reference databases (UniRef30 2021-03^109^, MGnify 2022-05^115^, BFD^114^ and Uniclust30^116^), with a maximum PDB template release date of 2020-05-14. Five structural models were generated, followed by GPU-Amber relaxation. The top-ranked model, selected based on the highest average pLDDT confidence score (pLDDT ≥ 90), was retained and compared using Foldseek against AFDB to infer its potential function.

### Statistical analysis and data visualization

All statistical analyses were performed in R (v4.5.0) using the specific tests described in the corresponding Methods sections. When multiple comparisons were conducted, P-values were adjusted for FDR using the Benjamini–Hochberg procedure, implemented via the p.adjust function in base R. Data visualization and figure generation were performed in R using ggplot2^117^ (v3.5.2) and BioRender. Protein structures in Figure 3J were analyzed with PyMOL^118^ (https://www.pymol.org/) and rendered with Mol* Viewer^119^. Visualization of the protein structure, VRs and amino acid positions of the DGR target in Figure 5F was performed with ChimeraX^120^ (v1.10.1).

## Data availability

Sample information, timepoint metadata, clinical data, family structure and quality-trimmed, human-decontaminated sequencing reads are available in the European Genome-Phenome Archive (EGA) as EGAS50000000133 and can be accessed upon submitting a request via the form at https://forms.gle/A4Jem2rMnjcygWRD6. Access to Lifelines NEXT Project data is granted to qualified researchers in accordance with the LLNEXT Data Access Agreement: https://groningenmicrobiome.org/?page_id=2598. This procedure ensures that data are used solely for research purposes and in compliance with the informed consent provided by Lifelines NEXT participants. To facilitate data access and usage, a comprehensive instruction guide is available at: https://github.com/GRONINGEN-MICROBIOME-CENTRE/LLNEXT_pilot/blob/main/Data_Access_EGA.md. The viral catalogs generated in this study are available at Zenodo (https://doi.org/10.5281/zenodo.18981021).

## Code availability

All scripts used for analysis and figure generation are publicly available at Zenodo (https://doi.org/10.5281/zenodo.18866521) and on GitHub at: https://github.com/GRONINGEN-MICROBIOME-CENTRE/Lifelines_NEXT_Virome_Penta.

## Acknowledgements

The data used in this manuscript were provided by LLNEXT. The authors are grateful to all the parents and infants who participated in LLNEXT and thank all staff involved in participant recruitment, maternal sample collection and the building and maintenance of the cohort. We also thank Kate Mc Intyre for editing the manuscript. We thank the UMCG Genomics Coordination Center and the Center for Information Technology of the University of Groningen and for their support and for providing access to the Gearshift, Nibbler and Hábrók high performance computing clusters. The Lifelines NEXT cohort study received funds from the University Medical Center Groningen Hereditary Metabolic Diseases Fund, Health~Holland (Top Sector Life Sciences and Health), the EU, the Northern Netherlands Alliance (SNN), the provinces of Friesland and Groningen, the municipality of Groningen, Philips, and the Société des Produits Nestlé. A.F.P. holds a De Cocks-Hadders Stichting grant (2023-50). T.S. holds a scholarship from the Junior Scientific Masterclass, University of Groningen and a De Cocks-Hadders Stichting grant (Winston Bakker Fonds WB-08). S.B. is supported by EUCAN-connect, a federated FAIR platform enabling large-scale analysis of high-value cohort data connecting Europe and Canada in personalized health, which received funding from the EU’s Horizon 2020 research innovation program (824989). J.E.S. holds a de Cock-Hadders Stichting grant (2021-57). The work of A.P.C and S.R. is conducted at the US Department of Energy Joint Genome Institute (https://ror.org/04xm1d337), a DOE Office of Science User Facility, supported by the Office of Science of the US Department of Energy operated under contract no. DE-AC02-05CH11231. J.F. is supported by the Dutch Heart Foundation IN-CONTROL (CVON2018-27), a European Research Council (ERC) Consolidator grant (grant agreement No. 101001678), Nederlandse Organisatie voor Wetenschappelijk Onderzoek (NWO) VICI grant VI.C.202.022, the AMMODO Science Award 2023 for Biomedical Sciences from Stichting Ammodo and NWO KIC grant KICH1.LWV04.21.01 (together with A.Z.). A.K. was supported by the NWO Gravitation grant Exposome-NL (024.004.017, together with A.Z.). S.G. was supported by a scholarship from the Graduate School of Medical Sciences, University of Groningen, a de Cock-Hadders Stitching grant (2021-08) and a NWO Rubicon grant (019.243EN.053). A.Z. was supported by the ERC Starting Grant 715772, NWO-VIDI grant 016.178.056, NWO-VICI grant VI.C.232.074, EU Horizon Europe Program grant INITIALISE (101094099), NWO KIC grant KICH1.LWV04.21.01 (together with J.F.) and NWO Gravitation grant Exposome-NL (024.004.017, together with A.K.).

## Author contributions

A.F.P., T.S., S.B., J.E.S., M.B.G. and S.G. processed the phenotypic data. T.S., S.B., J.E.S., S.G. and L.L.N. gathered and processed the biological data. A.F.P., T.S., S.B., J.E.S., M.B.G., A.K. and S.G. processed the metagenomic data. A.F.P., A.G., S.R. and S.G. developed the assembly-based viral discovery pipeline. A.F.P., A.G., N.K., and S.G. performed the data analysis. A.F.P., A.Gou. and A.R.M. performed structural analyses and visualization. A.G., A.P.C., J.F., A.K., S.R., S.G. and A.Z. supported and discussed the data analysis. A.F.P. performed the statistical analysis under the supervision of A.K. A.F.P. wrote the manuscript and created the figures. T.S., S.B., J.E.S., M.B.G., A.K., S.G., A.Z. and L.L.N. initiated, designed and supported the LLNEXT cohort study. All authors have read and agreed to the published version of the manuscript.

## Supplementary Information

### Lifelines NEXT cohort study authors and affiliations

Milla F. Brandão-Gois^1^, Marcel Bruinenberg^2^, Siobhan Brushett^1,3^, Jackie A.M. Dekens^1,4^, Sanzhima Garmaeva^1^, Sanne J. Gordijn^5^, Soesma A. Jankipersadsing^1^, Ank de Jonge^6,7^, Trynke R. de Jong^2^, Gerard H. Koppelman^8,9^, Marlou L.A de Kroon^3^, Folkert Kuipers^8,10^, Alexander M. Kurilshikov^1^, Lilian L. Peters^6,7^, Jelmer R. Prins^5^, Sijmen A. Reijneveld^3^, Sicco Scherjon^5^, Jan Sikkema^4^, Trishla Sinha^1^, Johanne E. Spreckels^1^, Aline B. Sprikkelman^8,9^, Morris A. Swertz^1^, Henkjan J. Verkade^8^, Cisca Wijmenga^1^, Alexandra Zhernakova^1^

^1^Department of Genetics, University of Groningen and University Medical Center Groningen, Groningen, the Netherlands

^2^Lifelines Cohort Study and Biobank, Groningen, the Netherlands

^3^Department of Health Sciences, University of Groningen and University Medical Center Groningen, Groningen, the Netherlands

^4^Innovation Center, University Medical Center Groningen, Groningen, the Netherlands

^5^Department of Obstetrics and Gynecology, University of Groningen and University Medical Center Groningen, Groningen, the Netherlands

^6^Department of Primary and Long-term Care, University of Groningen, University Medical Center Groningen, Groningen, the Netherlands

^7^Midwifery Science, Amsterdam University Medical Center, Vrije Universiteit Amsterdam, AVAG, Amsterdam Public Health, Amsterdam, the Netherlands

^8^Department of Pediatrics, University of Groningen and University Medical Center Groningen, Groningen, the Netherlands

^9^Groningen Research Institute for Asthma and COPD (GRIAC), University of Groningen and University Medical Center Groningen, Groningen, the Netherlands

^10^European Research Institute for the Biology of Ageing (ERIBA), University of Groningen and University Medical Center Groningen, Groningen, the Netherlands

---

**Extended Data Fig. 1.**
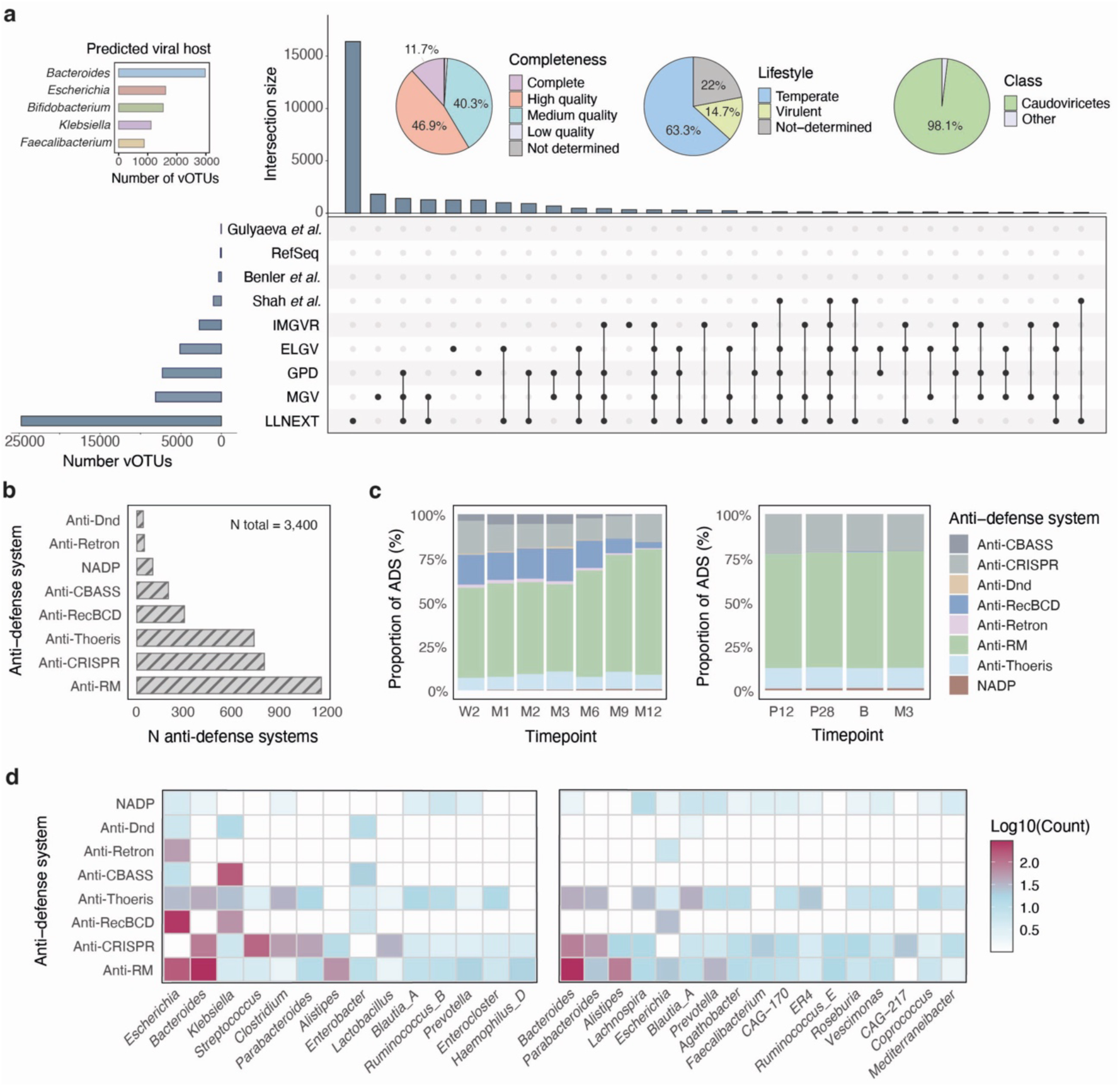
Characterization of the Lifelines NEXT maternal–infant gut DNA viral catalog (LLNEXT-GV) and its anti-defense system repertoire. **(A)** UpSet plot comparing the overlap of the LLNEXT-GV catalog (n = 31,019 vOTUs) and the public human gut virus databases included in the study: MGV (n = 189,680), GPD (n = 82,621), IMG/VR (n = 36,064), ELGV (n = 21,295), RefSeq (n = 5,199), Shah et al. (n = 4,627), Benler et al. (n = 1,480) and Gulyaeva et al. (n = 637). The Yutin et al. and Guerin et al. datasets are excluded from the plot due to their very limited representation. Intersection sizes represent the number of vOTUs containing at least one viral genome from each database combination (top 30 shown). Above, a bar plot and pie charts summarize the distribution of vOTUs by predicted viral host, estimated genome completeness, lifestyle and class-level taxonomy. **(B)** Bar plot showing the number of viral ADS detected across all LLNEXT-GV vOTUs. **(C)** Proportion of ADS in infants (left) and mothers (right) at each time point, estimated from the presence of vOTUs carrying each ADS type. **(D)** Heatmap showing the total number (log scale) of ADS detected in vOTUs according to their predicted bacterial host at the genus level for infants (left) and mothers (right). Only hosts with at least 20 ADS detected across their predicted phages are included. In all ADS-related plots, only the eight most common ADS types across all vOTUs are shown.

**Extended Data Fig. 2.**
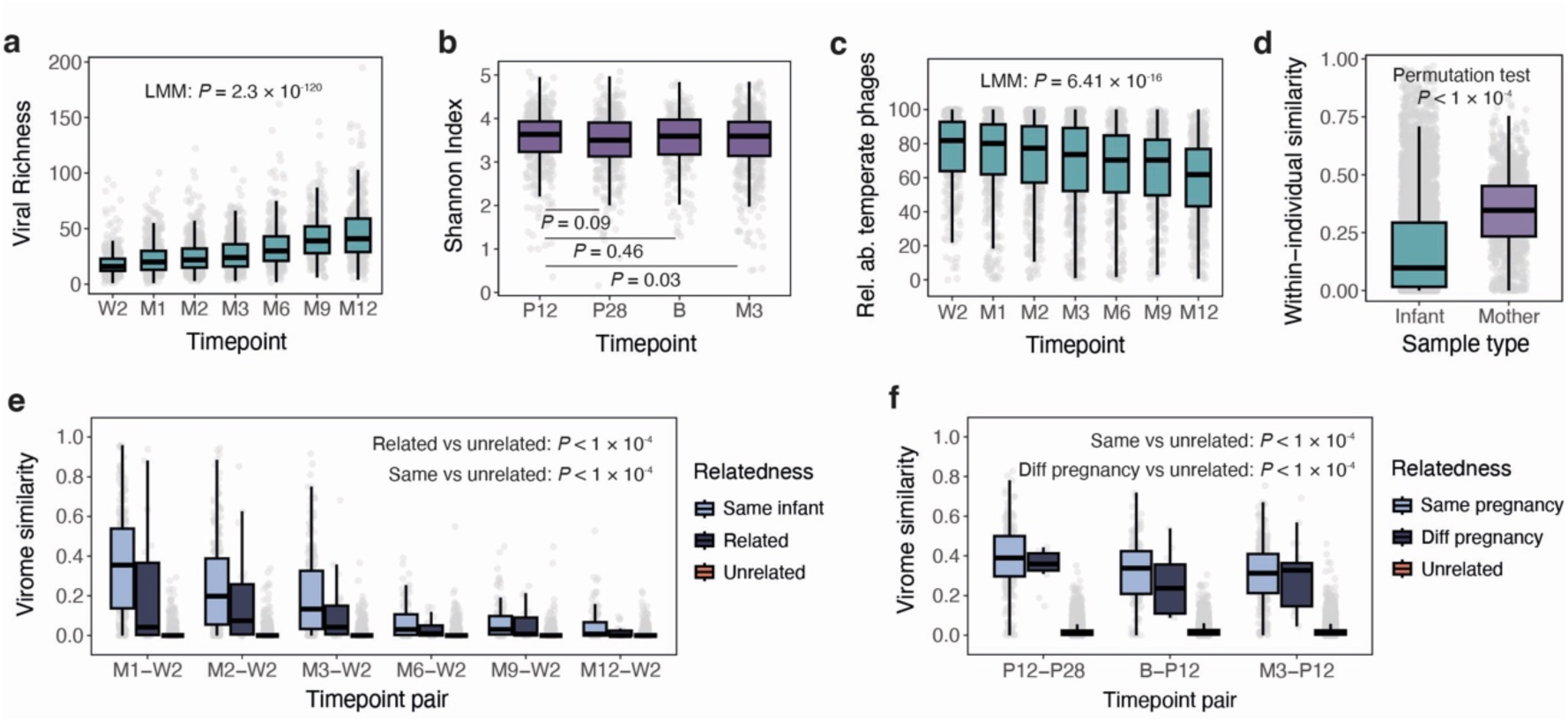
Virome dynamics across infants and mothers, with community similarity reflecting biological-relatedness. (**A**) Viral richness across infant timepoints, showing a significant increase over the first year of life. (**B**) Shannon diversity across maternal timepoints, showing only minor differences over time. (**C**) Longitudinal change in the relative abundance of temperate phages over time in infants. (**D**) Within-individual similarity of viral communities based on Bray-Curtis distances for mothers and infants. (**E**) Viral community similarity across infant timepoint pairs (comparing each timepoint to W2), stratified by whether samples originated from the same infant, related infants or unrelated infants. (**F**) Viral community similarity across maternal timepoint pairs (comparing each timepoint to P12), stratified by samples from the same pregnancy, different pregnancies or unrelated mothers. Boxplots show median, IQR, and 1.5 × IQR (whiskers). Gray points represent individual samples.

**Extended Data Fig. 3.**
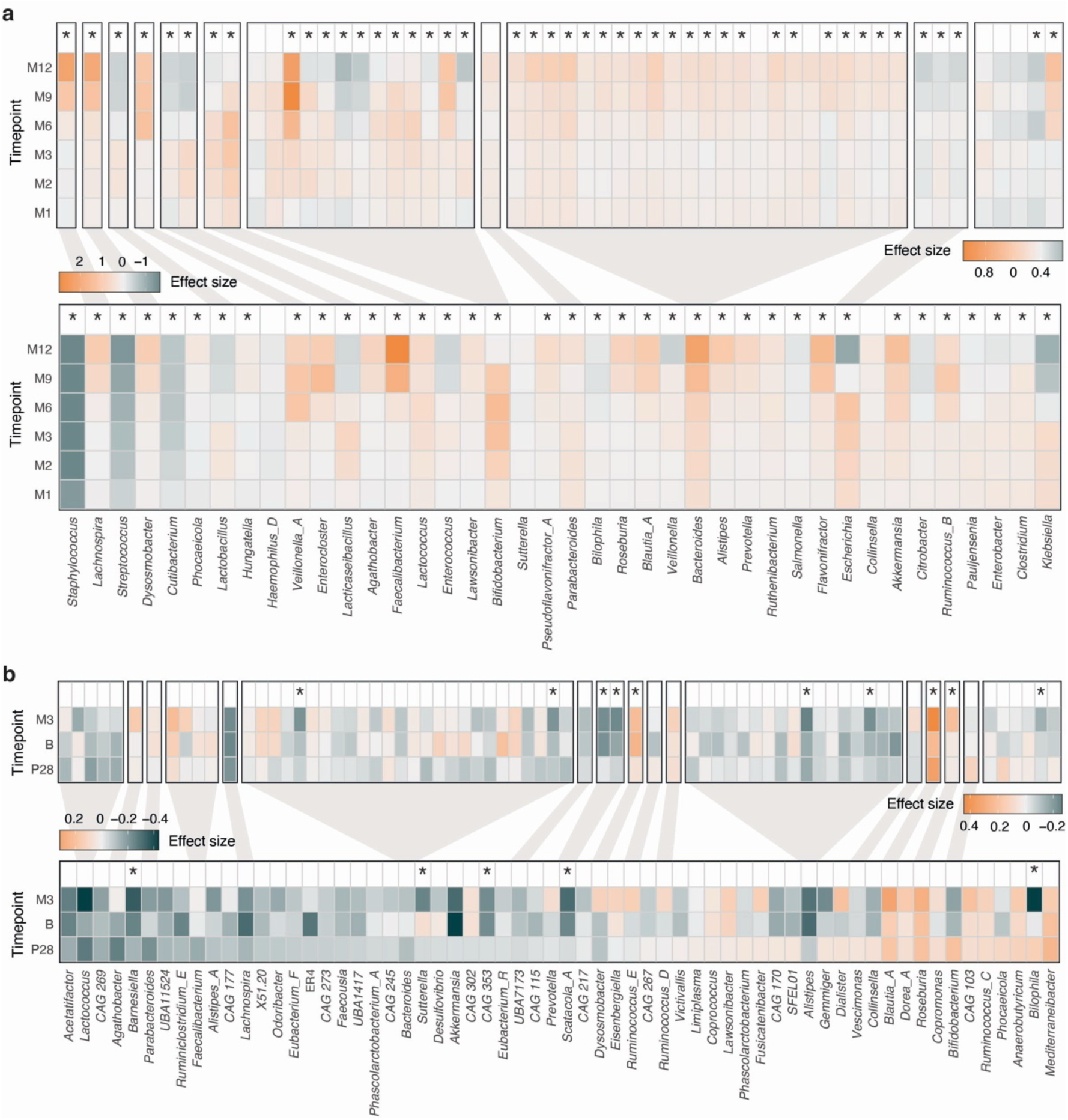
Temporal dynamics of vOTU- and host-level phage abundances in infants and mothers. **(A)** Heatmaps showing temporal variation in the abundance of prevalent vOTUs (top) and vOTUs grouped by their predicted bacterial host genus (bottom) across infant timepoints, with all comparisons performed against W2. (**B**) Analogous heatmaps for mothers, with all comparisons performed against P12. For both panels, associations between abundance and time were assessed using linear mixed models (LMMs) and only prevalent features (≥ 5% in infants, ≥ 10% in mothers) were included. Color scales represent effect sizes from the LMMs. Gray shades link individual vOTUs to their predicted host genus. * FDR < 0.05.

**Extended Data Fig. 4.**
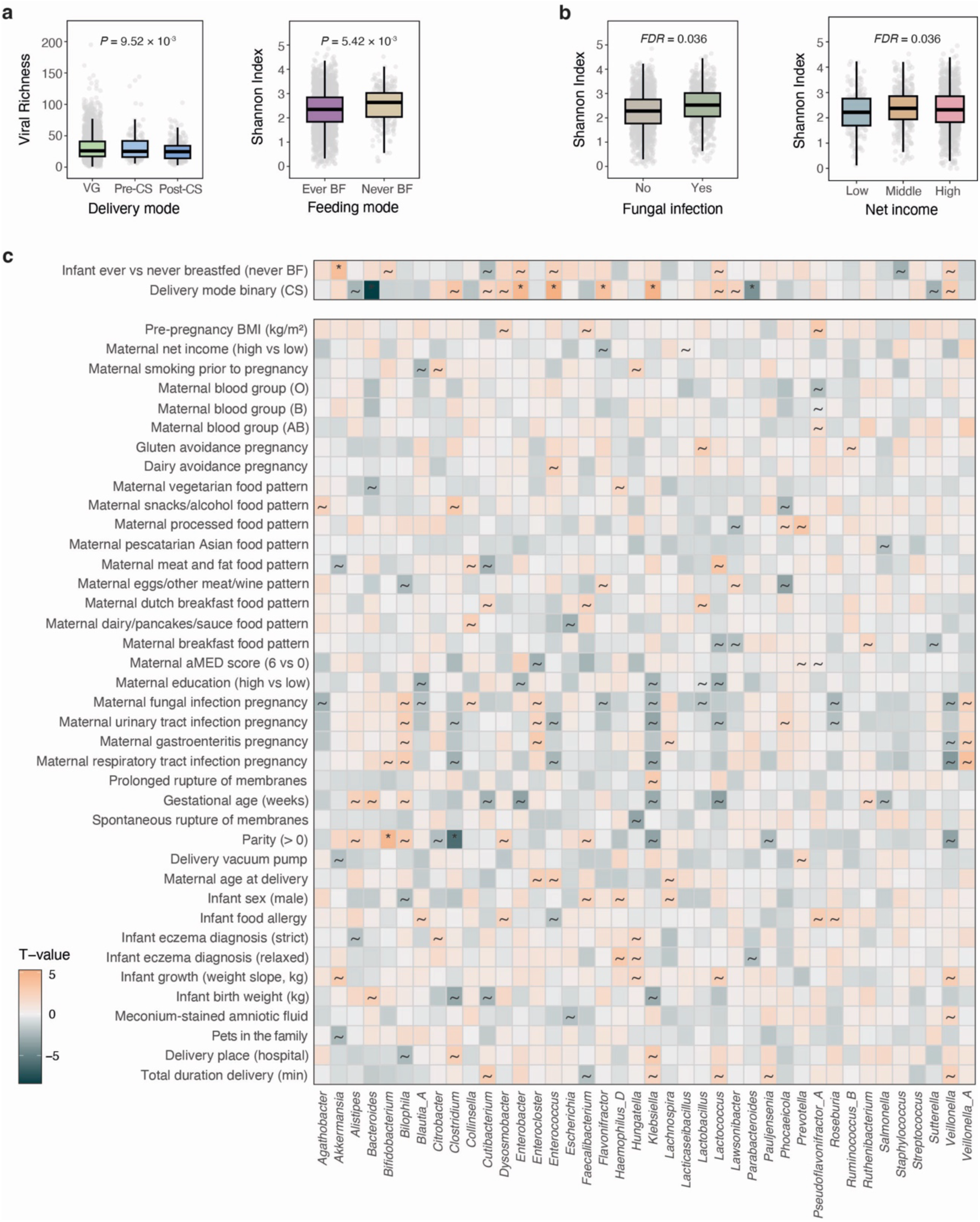
Infant gut virome features associated with host, delivery and feeding factors. **(A)** Infant viral richness stratified by delivery mode (left: vaginal (VG), pre-labor cesarean section (CS), post-labor CS) and feeding mode (right: ever vs. never breastfed). **(B)** Infant viral diversity (measured by Shannon Index) according to the presence of pregnancy vaginal fungal infections (left) and maternal net income (right). Nominally significant *P*-values are shown in (A). FDR-adjusted *P*-values accounting for bacterial diversity are shown in (B). Boxplots show the median, IQR and 1.5 × IQR whiskers. Individual points represent samples. **(C)** Heatmap of associations between viral abundances, grouped by predicted bacterial host and maternal and infant phenotypes. Associations were assessed using linear mixed models (LMMs), including read depth, DNA concentration and timepoint as fixed effects, with infant feeding mode and delivery mode added as additional covariates for all associations except those for those two phenotypes (shown on top). Only prevalent viral groups (≥ 5%) were included. Each cell represents the t-value from the linear model, with positive values in orange and negative values in dark gray. Only phenotypes with any nominally significant associations are highlighted. **~** nominally significant association. * FDR < 0.05.

**Extended Data Fig. 5.**
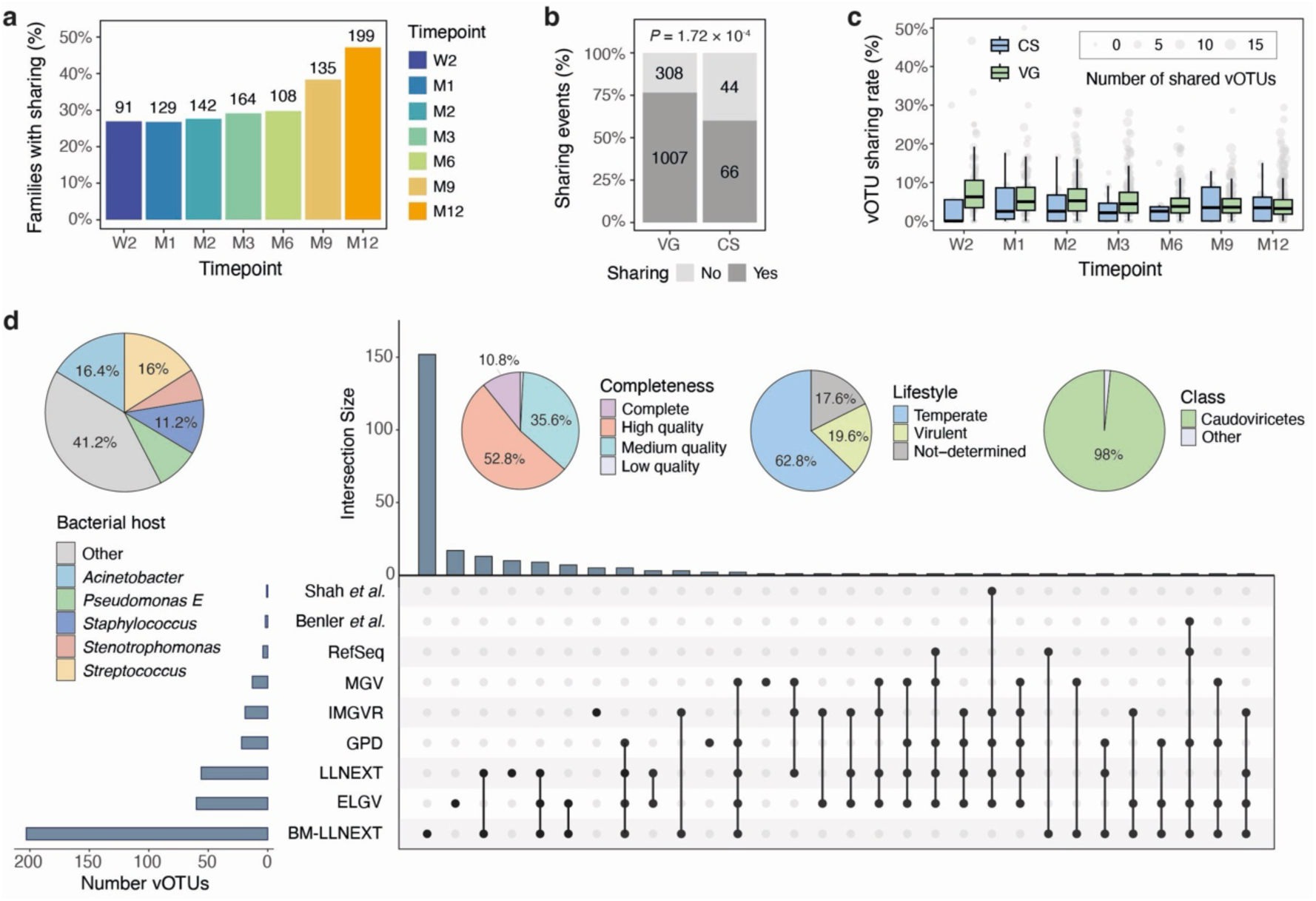
Viral strain-sharing between mothers and infants and characterization of the breastmilk virome. **(A)** Proportion of families with viral strain-sharing events at each infant timepoint (compared with any maternal timepoint). The number above each bar indicates the total number of families in which strain-sharing was detected. **(B)** Proportion of viral strain-sharing events stratified by delivery mode (VG: vaginal, CS: C-section). The total number of pairwise strain comparisons classified as sharing events (popANI >99.999%, dark gray) or non-sharing (popANI ≤ 99.999%, light gray) is indicated in each bar. (**C**) vOTU sharing rate (number of shared vOTUs/total number of infant vOTUs) across infant timepoints, stratified by delivery mode. Boxplots show the median, IQR and 1.5 × IQR whiskers. Individual gray points represent samples. **(D)** UpSet plot comparing the overlap of breastmilk vOTUs with several public human gut virus databases and the LLNEXT-GV catalog. The datasets from Gulyaeva et al., Yutin et al. and Guerin et al. were excluded from the plot due to their very limited representation. Intersection sizes represent the number of vOTUs containing at least one viral genome from each database combination (top 30 shown). Pie charts summarize the distribution of vOTUs by predicted bacterial host, estimated genome completeness, lifestyle and class-level taxonomy.

**Extended Data Fig. 6.**
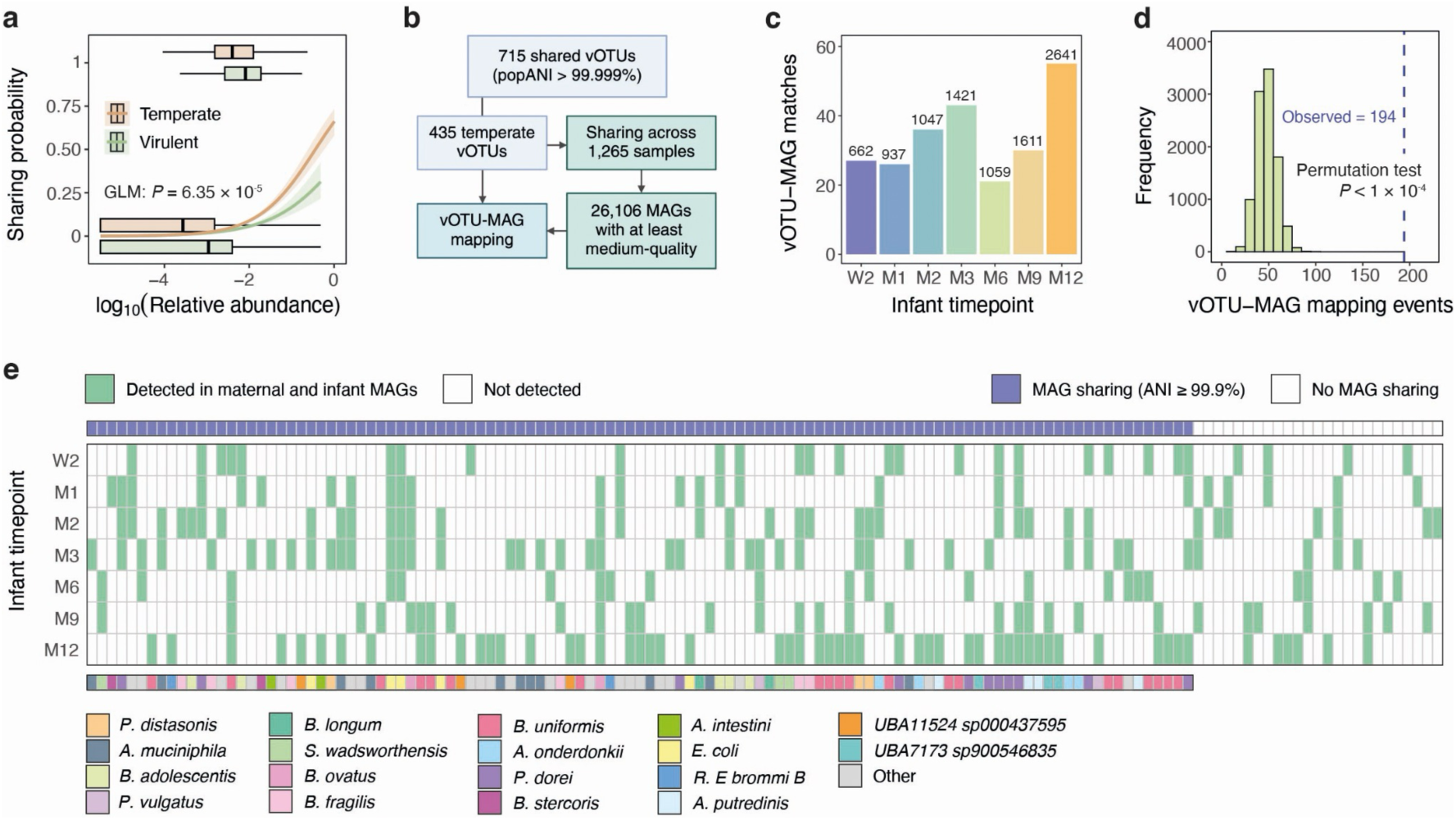
Co-transmission of temperate viruses and their bacterial hosts. **(A)** Relationship between vOTU mean abundance in mothers and the probability of sharing, shown separately for temperate and virulent lifestyles. Logistic regression curves show the modeled probability of persistence across the range of log10-transformed mean abundances, with shaded areas representing 95% confidence intervals. Boxplots show the distribution of mean abundances for shared (top) and non-shared (bottom) vOTUs within each lifestyle category. **(B)** Workflow for mapping potentially shared temperate vOTUs to medium and high-quality metagenome-assembled genomes (MAGs) reconstructed from the maternal and infant samples where viral strain-sharing was detected. (**C**) Bar plot showing the number of vOTU‒MAG matching events (where the vOTU mapped to a MAG in both the mother and infant) at each infant timepoint. The total number of MAGs included from infants at each timepoint is indicated above each bar. (**D**) Histogram comparing the frequency of vOTU‒MAG matching events observed in related mother–infant pairs (Observed) to the frequency expected from randomly paired mother–infant samples (Permutation test, p < 1 × 10^−4^). Dashed blue line indicates the observed value. **(E)** Heatmap of vOTU‒MAG matching and strain-sharing events. Each column represents a potential sharing event. Green indicates that the shared temperate vOTU was detected in both maternal and infant MAGs. Blue in the top bar indicates whether the corresponding maternal and infant MAGs were also shared strains (ANI >99.9%). The color bar at the bottom indicates the species of the matched MAG for cases where MAG-sharing was detected.

**Extended Data Fig. 7.**
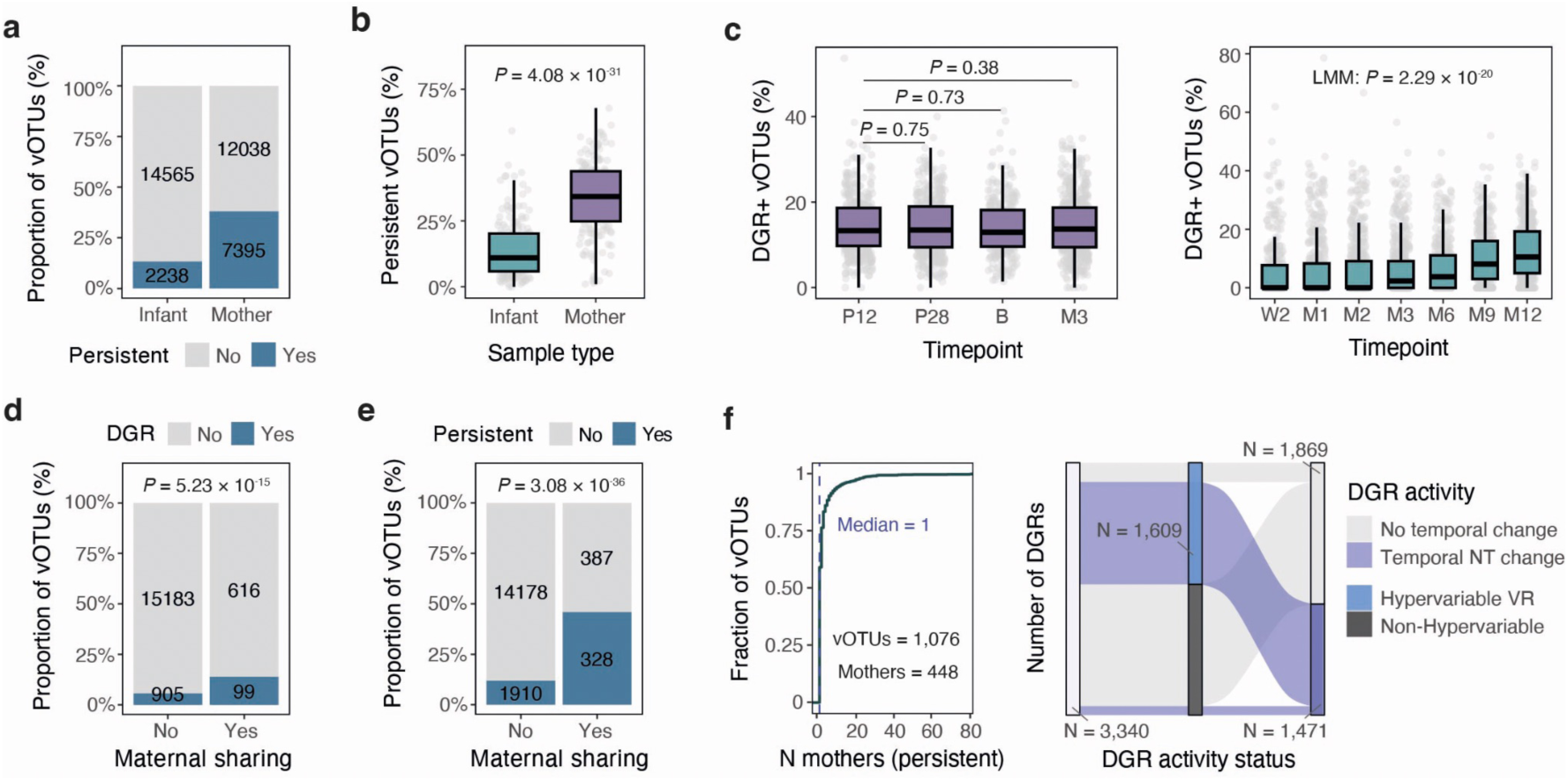
Viral persistence and DGR characterization in mothers and infants. **(A)** Proportion of infant and maternal vOTUs identified as persistent (Methods). **(B)** Comparison of the proportion of persistent vOTUs per individual within a subset of infants (n = 166) and mothers (n = 174) with near-complete longitudinal sampling. (**C**) Proportion of DGR-encoding vOTUs (DGR+) across mothers (left) and infants (right), per timepoint. Boxplots show median, IQR and 1.5 × IQR whiskers. Gray points represent individual samples. **(D–E)** Proportion of (**D**) DGR+ vOTUs and (**E**) persistent vOTUs stratified by maternal sharing status. Numbers within bars indicate vOTU counts. (**F**) At left, the cumulative distribution of the number of mothers in which individual vOTUs persist over time. The dashed line indicates the median number of mothers (median = 1). At right, an alluvial plot summarizes DGR activity status in mothers, classified according to the presence of higher nucleotide variability in VR vs non-VR regions and the detection of temporal nucleotide changes in the VR region.

## References

1 Tanaka, M. & Nakayama, J. Development of the gut microbiota in infancy and its impact on health in later life. Allergol Int 66, 515–522 (2017). 10.1016/j.alit.2017.07.010

2 Jian, C. et al. Early-life gut microbiota and its connection to metabolic health in children: Perspective on ecological drivers and need for quantitative approach. EBioMedicine 69, 103475 (2021). 10.1016/j.ebiom.2021.103475

3 Sanidad, K. Z. & Zeng, M. Y. Neonatal gut microbiome and immunity. Curr Opin Microbiol 56, 30–37 (2020). 10.1016/j.mib.2020.05.011

4 Tamburini, S., Shen, N., Wu, H. C. & Clemente, J. C. The microbiome in early life: implications for health outcomes. Nat Med 22, 713–722 (2016). 10.1038/nm.4142

5 Enav, H., Bäckhed, F. & Ley, R. E. The developing infant gut microbiome: A strain-level view. Cell Host Microbe 30, 627–638 (2022). 10.1016/j.chom.2022.04.009

6 Bogaert, D. et al. Mother-to-infant microbiota transmission and infant microbiota development across multiple body sites. Cell Host Microbe 31, 447–460.e446 (2023). 10.1016/j.chom.2023.01.018

7 Sinha, T. et al. Pregnancy and Early Life Gut Microbiome: Influencing Factors and Health Implications. Research Square (2024).

8 Stewart, C. J. et al. Temporal development of the gut microbiome in early childhood from the TEDDY study. Nature 562, 583–588 (2018). 10.1038/s41586-018-0617-x

9 Roswall, J. et al. Developmental trajectory of the healthy human gut microbiota during the first 5 years of life. Cell Host Microbe 29, 765–776.e763 (2021). 10.1016/j.chom.2021.02.021

10 Bokulich, N. A. et al. Antibiotics, birth mode, and diet shape microbiome maturation during early life. Sci Transl Med 8, 343ra382 (2016). 10.1126/scitranslmed.aad7121

11 Reyman, M. et al. Impact of delivery mode-associated gut microbiota dynamics on health in the first year of life. Nat Commun 10, 4997 (2019). 10.1038/s41467-019-13014-7

12 Bäckhed, F. et al. Dynamics and Stabilization of the Human Gut Microbiome during the First Year of Life. Cell Host Microbe 17, 690–703 (2015). 10.1016/j.chom.2015.04.004

13 Cao, Z. et al. The gut virome: A new microbiome component in health and disease. EBioMedicine 81, 104113 (2022). 10.1016/j.ebiom.2022.104113

14 Ogilvie, L. A. & Jones, B. V. The human gut virome: a multifaceted majority. Front Microbiol 6, 918 (2015). 10.3389/fmicb.2015.00918

15 Chica Cardenas, L. A., Leonard, M. M., Baldridge, M. T. & Handley, S. A. Gut virome dynamics: from commensal to critical player in health and disease. Nat Rev Gastroenterol Hepatol 23, 126–144 (2026). 10.1038/s41575-025-01134-z

16 Borin, J. M. et al. Fecal virome transplantation is sufficient to alter fecal microbiota and drive lean and obese body phenotypes in mice. Gut Microbes 15, 2236750 (2023). 10.1080/19490976.2023.2236750

17 Roux, S. et al. Minimum Information about an Uncultivated Virus Genome (MIUViG). Nat Biotechnol 37, 29–37 (2019). 10.1038/nbt.4306

18 Garmaeva, S. et al. Studying the gut virome in the metagenomic era: challenges and perspectives. BMC Biol 17, 84 (2019). 10.1186/s12915-019-0704-y

19 Leal Rodríguez, C., et al. The infant gut virome is associated with preschool asthma risk independently of bacteria. Nat Med 30, 138–148 (2024). 10.1038/s41591-023-02685-x

20 Perdue, T. J. et al. The enteric DNA virome differs in infants at risk for atopic disease. Gut Microbes 18, 2616066 (2026). 10.1080/19490976.2026.2616066

21 Vehik, K. et al. Prospective virome analyses in young children at increased genetic risk for type 1 diabetes. Nat Med 25, 1865–1872 (2019). 10.1038/s41591-019-0667-0

22 Garmaeva, S. et al. Transmission and dynamics of mother-infant gut viruses during pregnancy and early life. Nat Commun 15, 1945 (2024). 10.1038/s41467-024-45257-4

23 Fernández-Pato, A. et al. Early-life development of the gut virome and plasmidome: A longitudinal study in cesarean-born infants. Cell Rep 44, 115731 (2025). 10.1016/j.celrep.2025.115731

24 Duranti, S. et al. Maternal inheritance of bifidobacterial communities and bifidophages in infants through vertical transmission. Microbiome 5, 66 (2017). 10.1186/s40168-017-0282-6

25 Pannaraj, P. S. et al. Shared and Distinct Features of Human Milk and Infant Stool Viromes. Front Microbiol 9, 1162 (2018). 10.3389/fmicb.2018.01162

26 Maqsood, R. et al. Discordant transmission of bacteria and viruses from mothers to babies at birth. Microbiome 7, 156 (2019). 10.1186/s40168-019-0766-7

27 Liang, G. et al. The stepwise assembly of the neonatal virome is modulated by breastfeeding. Nature 581, 470–474 (2020). 10.1038/s41586-020-2192-1

28 Guo, Y. et al. Phage diversity in human breast milk: a systematic review. Eur J Pediatr 184, 334 (2025). 10.1007/s00431-025-06173-x

29 Lourenço, M. et al. The Spatial Heterogeneity of the Gut Limits Predation and Fosters Coexistence of Bacteria and Bacteriophages. Cell Host Microbe 28, 390–401.e395 (2020). 10.1016/j.chom.2020.06.002

30 Chin, W. H. et al. Bacteriophages evolve enhanced persistence to a mucosal surface. Proc Natl Acad Sci U S A 119, e2116197119 (2022). 10.1073/pnas.2116197119

31 Popescu, M., Van Belleghem, J. D., Khosravi, A. & Bollyky, P. L. Bacteriophages and the Immune System. Annu Rev Virol 8, 415–435 (2021). 10.1146/annurev-virology-091919-074551

32 Lou, Y. C. et al. Infant gut DNA bacteriophage strain persistence during the first 3 years of life. Cell Host Microbe 32, 35–47.e36 (2024). 10.1016/j.chom.2023.11.015

33 De Sordi, L., Lourenço, M. & Debarbieux, L. “I will survive”: A tale of bacteriophage-bacteria coevolution in the gut. Gut Microbes 10, 92–99 (2019). 10.1080/19490976.2018.1474322

34 Shkoporov, A. N. et al. The Human Gut Virome Is Highly Diverse, Stable, and Individual Specific. Cell Host Microbe 26, 527–541.e525 (2019). 10.1016/j.chom.2019.09.009

35 Murtazalieva, K., Mu, A., Petrovskaya, A. & Finn, R. D. The growing repertoire of phage anti-defence systems. Trends Microbiol 32, 1212–1228 (2024). 10.1016/j.tim.2024.05.005

36 Niault, T., van Houte, S., Westra, E. & Swarts, D. C. Evolution and ecology of anti-defence systems in phages and plasmids. Curr Biol 35, R32–r44 (2025). 10.1016/j.cub.2024.11.033

37. Camargo, A. P., et al. A genomic atlas of the human gut virome elucidates genetic factors shaping host interactions. bioRxiv (2025).

38 Warmink-Perdijk, W. D. B. et al. Lifelines NEXT: a prospective birth cohort adding the next generation to the three-generation Lifelines cohort study. Eur J Epidemiol 35, 157–168 (2020). 10.1007/s10654-020-00614-7

39 Shenhav, L. et al. Microbial colonization programs are structured by breastfeeding and guide healthy respiratory development. Cell 187, 5431–5452.e5420 (2024). 10.1016/j.cell.2024.07.022

40 Mitchell, C. M. et al. Delivery Mode Affects Stability of Early Infant Gut Microbiota. Cell Rep Med 1, 100156 (2020). 10.1016/j.xcrm.2020.100156

41 Shi, P. et al. TEMPTED: time-informed dimensionality reduction for longitudinal microbiome studies. Genome Biol 25, 317 (2024). 10.1186/s13059-024-03453-x

42 Ferretti, P. et al. Mother-to-Infant Microbial Transmission from Different Body Sites Shapes the Developing Infant Gut Microbiome. Cell Host Microbe 24, 133–145.e135 (2018). 10.1016/j.chom.2018.06.005

43 Yassour, M. et al. Strain-Level Analysis of Mother-to-Child Bacterial Transmission during the First Few Months of Life. Cell Host Microbe 24, 146–154.e144 (2018). 10.1016/j.chom.2018.06.007

44 Salter, S. J. et al. Reagent and laboratory contamination can critically impact sequence-based microbiome analyses. BMC Biol 12, 87 (2014). 10.1186/s12915-014-0087-z

45 Spreckels, J. E. et al. Host and environmental determinants of human milk oligosaccharides and microbiota in the Lifelines NEXT cohort. Cell Rep 44, 116124 (2025). 10.1016/j.celrep.2025.116124

46 Notarbartolo, V., Giuffrè, M., Montante, C., Corsello, G. & Carta, M. Composition of Human Breast Milk Microbiota and Its Role in Children’s Health. Pediatr Gastroenterol Hepatol Nutr 25, 194–210 (2022). 10.5223/pghn.2022.25.3.194

47 García-López, M. et al. Analysis of 1,000 Type-Strain Genomes Improves Taxonomic Classification of Bacteroidetes. Front Microbiol 10, 2083 (2019). 10.3389/fmicb.2019.02083

48 Wirbel, J. et al. Long-read metagenomics reveals phage dynamics in the human gut microbiome. Nature 649, 982–990 (2026). 10.1038/s41586-025-09786-2

49 Bignaud, A. et al. Phages with a broad host range are common across ecosystems. Nat Microbiol 10, 2537–2549 (2025). 10.1038/s41564-025-02108-2

50 Terzian, P. et al. PHROG: families of prokaryotic virus proteins clustered using remote homology. NAR Genom Bioinform 3, lqab067 (2021). 10.1093/nargab/lqab067

51 Aramaki, T. et al. KofamKOALA: KEGG Ortholog assignment based on profile HMM and adaptive score threshold. Bioinformatics 36, 2251–2252 (2020). 10.1093/bioinformatics/btz859

52 Tesson, F. et al. Exploring the diversity of anti-defense systems across prokaryotes, phages and mobile genetic elements. Nucleic Acids Res 53 (2025). 10.1093/nar/gkae1171

53 Mirdita, M. et al. ColabFold: making protein folding accessible to all. Nat Methods 19, 679–682 (2022). 10.1038/s41592-022-01488-1

54 Berman, H. M. et al. The Protein Data Bank. Nucleic Acids Res 28, 235–242 (2000). 10.1093/nar/28.1.235

55 Varadi, M. et al. AlphaFold Protein Structure Database in 2024: providing structure coverage for over 214 million protein sequences. Nucleic Acids Res 52, D368–d375 (2024). 10.1093/nar/gkad1011

56 Kim, R. S., Levy Karin, E., Mirdita, M., Chikhi, R. & Steinegger, M. BFVD-a large repository of predicted viral protein structures. Nucleic Acids Res 53, D340–d347 (2025). 10.1093/nar/gkae1119

57 van Kempen, M. et al. Fast and accurate protein structure search with Foldseek. Nat Biotechnol 42, 243–246 (2024). 10.1038/s41587-023-01773-0

58 Hör, J., Wolf, S. G. & Sorek, R. Bacteria conjugate ubiquitin-like proteins to interfere with phage assembly. Nature 631, 850–856 (2024). 10.1038/s41586-024-07616-5

59 Chambers, L. R. et al. A eukaryotic-like ubiquitination system in bacterial antiviral defence. Nature 631, 843–849 (2024). 10.1038/s41586-024-07730-4

60 Guo, X. et al. Structure and mechanism of a phage-encoded SAM lyase revises catalytic function of enzyme family. Elife 10 (2021). 10.7554/eLife.61818

61 Chi, H. et al. SAM-AMP lyases in type III CRISPR defence. Nucleic Acids Res 53 (2025). 10.1093/nar/gkaf655

62 Roux, S. et al. Ecology and molecular targets of hypermutation in the global microbiome. Nat Commun 12, 3076 (2021). 10.1038/s41467-021-23402-7

63 Macadangdang, B. R. et al. Targeted protein evolution in the gut microbiome by diversity-generating retroelements. Science 390, eadv2111 (2025). 10.1126/science.adv2111

64 Zeng, S. et al. A metagenomic catalog of the early-life human gut virome. Nat Commun 15, 1864 (2024). 10.1038/s41467-024-45793-z

65 Nayfach, S. et al. Metagenomic compendium of 189,680 DNA viruses from the human gut microbiome. Nat Microbiol 6, 960–970 (2021). 10.1038/s41564-021-00928-6

66 Camarillo-Guerrero, L. F., Almeida, A., Rangel-Pineros, G., Finn, R. D. & Lawley, T. D. Massive expansion of human gut bacteriophage diversity. Cell 184, 1098–1109.e1099 (2021). 10.1016/j.cell.2021.01.029

67 McCann, A. et al. Viromes of one year old infants reveal the impact of birth mode on microbiome diversity. PeerJ 6, e4694 (2018). 10.7717/peerj.4694

68 Azad, M. B. et al. Gut microbiota of healthy Canadian infants: profiles by mode of delivery and infant diet at 4 months. Cmaj 185, 385–394 (2013). 10.1503/cmaj.121189

69 Bergström, A. et al. Establishment of intestinal microbiota during early life: a longitudinal, explorative study of a large cohort of Danish infants. Appl Environ Microbiol 80, 2889–2900 (2014). 10.1128/aem.00342-14

70 Chen, D. W. & Garud, N. R. Rapid evolution and strain turnover in the infant gut microbiome. Genome Res 32, 1124–1136 (2022). 10.1101/gr.276306.121

71 Fahur Bottino, G., et al. Early life microbial succession in the gut follows common patterns in humans across the globe. Nat Commun 16, 660 (2025). 10.1038/s41467-025-56072-w

72 Drozdz, M., Piekarowicz, A., Bujnicki, J. M. & Radlinska, M. Novel non-specific DNA adenine methyltransferases. Nucleic Acids Res 40, 2119–2130 (2012). 10.1093/nar/gkr1039

73 Medhekar, B. & Miller, J. F. Diversity-generating retroelements. Curr Opin Microbiol 10, 388–395 (2007). 10.1016/j.mib.2007.06.004

74 Obeng, N., Pratama, A. A. & Elsas, J. D. V. The Significance of Mutualistic Phages for Bacterial Ecology and Evolution. Trends Microbiol 24, 440–449 (2016). 10.1016/j.tim.2015.12.009

75 Scholtens, S. et al. Cohort Profile: LifeLines, a three-generation cohort study and biobank. Int J Epidemiol 44, 1172–1180 (2015). 10.1093/ije/dyu229

76 Nurk, S., Meleshko, D., Korobeynikov, A. & Pevzner, P. A. metaSPAdes: a new versatile metagenomic assembler. Genome Res 27, 824–834 (2017). 10.1101/gr.213959.116

77 Guo, J. et al. VirSorter2: a multi-classifier, expert-guided approach to detect diverse DNA and RNA viruses. Microbiome 9, 37 (2021). 10.1186/s40168-020-00990-y

78 Ren, J. et al. Identifying viruses from metagenomic data using deep learning. Quant Biol 8, 64–77 (2020). 10.1007/s40484-019-0187-4

79 Camargo, A. P. et al. Identification of mobile genetic elements with geNomad. Nat Biotechnol 42, 1303–1312 (2024). 10.1038/s41587-023-01953-y

80 Chen, L. & Banfield, J. F. COBRA improves the completeness and contiguity of viral genomes assembled from metagenomes. Nat Microbiol 9, 737–750 (2024). 10.1038/s41564-023-01598-2

81 Nayfach, S. et al. CheckV assesses the quality and completeness of metagenome-assembled viral genomes. Nat Biotechnol 39, 578–585 (2021). 10.1038/s41587-020-00774-7

82 Camargo, A. P. et al. IMG/VR v4: an expanded database of uncultivated virus genomes within a framework of extensive functional, taxonomic, and ecological metadata. Nucleic Acids Res 51, D733–d743 (2023). 10.1093/nar/gkac1037

83 Goldfarb, T. et al. NCBI RefSeq: reference sequence standards through 25 years of curation and annotation. Nucleic Acids Res 53, D243–d257 (2025). 10.1093/nar/gkae1038

84 Shah, S. A. et al. Expanding known viral diversity in the healthy infant gut. Nat Microbiol 8, 986–998 (2023). 10.1038/s41564-023-01345-7

85 Benler, S. et al. Thousands of previously unknown phages discovered in whole-community human gut metagenomes. Microbiome 9, 78 (2021). 10.1186/s40168-021-01017-w

86 Gulyaeva, A. et al. Discovery, diversity, and functional associations of crAss-like phages in human gut metagenomes from four Dutch cohorts. Cell Rep 38, 110204 (2022). 10.1016/j.celrep.2021.110204

87 Yutin, N. et al. Analysis of metagenome-assembled viral genomes from the human gut reveals diverse putative CrAss-like phages with unique genomic features. Nat Commun 12, 1044 (2021). 10.1038/s41467-021-21350-w

88 Guerin, E. et al. Biology and Taxonomy of crAss-like Bacteriophages, the Most Abundant Virus in the Human Gut. Cell Host Microbe 24, 653–664.e656 (2018). 10.1016/j.chom.2018.10.002

89 Langmead, B. & Salzberg, S. L. Fast gapped-read alignment with Bowtie 2. Nat Methods 9, 357–359 (2012). 10.1038/nmeth.1923

90 Quinlan, A. R. & Hall, I. M. BEDTools: a flexible suite of utilities for comparing genomic features. Bioinformatics 26, 841–842 (2010). 10.1093/bioinformatics/btq033

91 Zheng, K. et al. VITAP: a high precision tool for DNA and RNA viral classification based on meta-omic data. Nat Commun 16, 2226 (2025). 10.1038/s41467-025-57500-7

92 Black, E. J. et al. Virus taxonomy: the database of the International Committee on Taxonomy of Viruses. Nucleic Acids Res 54, D776–d789 (2026). 10.1093/nar/gkaf1159

93 Hockenberry, A. J. & Wilke, C. O. BACPHLIP: predicting bacteriophage lifestyle from conserved protein domains. PeerJ 9, e11396 (2021). 10.7717/peerj.11396

94 Johansen, J. et al. Centenarians have a diverse gut virome with the potential to modulate metabolism and promote healthy lifespan. Nat Microbiol 8, 1064–1078 (2023). 10.1038/s41564-023-01370-6

95 Roux, S. et al. iPHoP: An integrated machine learning framework to maximize host prediction for metagenome-derived viruses of archaea and bacteria. PLoS Biol 21, e3002083 (2023). 10.1371/journal.pbio.3002083

96 Blanco-Míguez, A. et al. Extending and improving metagenomic taxonomic profiling with uncharacterized species using MetaPhlAn 4. Nat Biotechnol 41, 1633–1644 (2023). 10.1038/s41587-023-01688-w

97 Uritskiy, G. V., DiRuggiero, J. & Taylor, J. MetaWRAP-a flexible pipeline for genome-resolved metagenomic data analysis. Microbiome 6, 158 (2018). 10.1186/s40168-018-0541-1

98 Parks, D. H., Imelfort, M., Skennerton, C. T., Hugenholtz, P. & Tyson, G. W. CheckM: assessing the quality of microbial genomes recovered from isolates, single cells, and metagenomes. Genome Res 25, 1043–1055 (2015). 10.1101/gr.186072.114

99. vegan: Community Ecology Package v. 2.6-6.1 (2024).

100 Kuznetsova, A., Brockhoff, P. B. & Christensen, R. H. B. lmerTest Package: Tests in Linear Mixed Effects Models. Journal of Statistical Software 82, 1–26 (2017). 10.18637/jss.v082.i13

101 Olm, M. R. et al. inStrain profiles population microdiversity from metagenomic data and sensitively detects shared microbial strains. Nat Biotechnol 39, 727–736 (2021). 10.1038/s41587-020-00797-0

102 Kuhn, M. & Wickham, H. Tidymodels: a collection of packages for modeling and machine learning using tidyverse principles. (2020).

103 Robin, X. et al. pROC: an open-source package for R and S+ to analyze and compare ROC curves. BMC Bioinformatics 12, 77 (2011). 10.1186/1471-2105-12-77

104 Li, H. Minimap2: pairwise alignment for nucleotide sequences. Bioinformatics 34, 3094–3100 (2018). 10.1093/bioinformatics/bty191

105 Shaw, J. & Yu, Y. W. Fast and robust metagenomic sequence comparison through sparse chaining with skani. Nat Methods 20, 1661–1665 (2023). 10.1038/s41592-023-02018-3

106 Hyatt, D. et al. Prodigal: prokaryotic gene recognition and translation initiation site identification. BMC Bioinformatics 11, 119 (2010). 10.1186/1471-2105-11-119

107 Steinegger, M. & Söding, J. MMseqs2 enables sensitive protein sequence searching for the analysis of massive data sets. Nat Biotechnol 35, 1026–1028 (2017). 10.1038/nbt.3988

108 Finn, R. D., Clements, J. & Eddy, S. R. HMMER web server: interactive sequence similarity searching. Nucleic Acids Res 39, W29–37 (2011). 10.1093/nar/gkr367

109 Suzek, B. E., Wang, Y., Huang, H., McGarvey, P. B. & Wu, C. H. UniRef clusters: a comprehensive and scalable alternative for improving sequence similarity searches. Bioinformatics 31, 926–932 (2015). 10.1093/bioinformatics/btu739

110 Steinegger, M. et al. HH-suite3 for fast remote homology detection and deep protein annotation. BMC Bioinformatics 20, 473 (2019). 10.1186/s12859-019-3019-7

111 Brooks, M. E. et al. glmmTMB Balances Speed and Flexibility Among Packages for Zero-inflated Generalized Linear Mixed Modeling. The R Journal 9, 378--400 (2017). 10.32614/RJ-2017-066

112 Camacho, C. et al. BLAST+: architecture and applications. BMC Bioinformatics 10, 421 (2009). 10.1186/1471-2105-10-421

113 Eren, A. M. et al. Anvi’o: an advanced analysis and visualization platform for ‘omics data. PeerJ 3, e1319 (2015). 10.7717/peerj.1319

114 Jumper, J. et al. Highly accurate protein structure prediction with AlphaFold. Nature 596, 583–589 (2021). 10.1038/s41586-021-03819-2

115 Richardson, L. et al. MGnify: the microbiome sequence data analysis resource in 2023. Nucleic Acids Res 51, D753–d759 (2023). 10.1093/nar/gkac1080

116 Mirdita, M. et al. Uniclust databases of clustered and deeply annotated protein sequences and alignments. Nucleic Acids Res 45, D170–d176 (2017). 10.1093/nar/gkw1081

117 Wickham, H. ggplot2: Elegant Graphics for Data Analysis. (Springer-Verlag New York, 2016).

118 Rigsby, R. E. & Parker, A. B. Using the PyMOL application to reinforce visual understanding of protein structure. Biochem Mol Biol Educ 44, 433–437 (2016). 10.1002/bmb.20966

119 Sehnal, D. et al. Mol* Viewer: modern web app for 3D visualization and analysis of large biomolecular structures. Nucleic Acids Res 49, W431–w437 (2021). 10.1093/nar/gkab314

120 Pettersen, E. F. et al. UCSF ChimeraX: Structure visualization for researchers, educators, and developers. Protein Sci 30, 70–82 (2021). 10.1002/pro.3943

